# Growth disrupting mutations in epigenetic regulatory molecules are associated with abnormalities of epigenetic aging

**DOI:** 10.1101/477356

**Authors:** Aaron R Jeffries, Reza Maroofian, Claire G. Salter, Barry A. Chioza, Harold E. Cross, Michael A. Patton, I. Karen Temple, Deborah Mackay, Faisal I. Rezwan, Lise Aksglæde, Diana Baralle, Tabib Dabir, Matthew Frank Hunter, Arveen Kamath, Ajith Kumar, Ruth Newbury-Ecob, Angelo Selicorni, Amanda Springer, Lionel van Maldergem, Vinod Varghese, Naomi Yachelevich, Katrina Tatton-Brown, Jonathan Mill, Andrew H. Crosby, Emma Baple

## Abstract

Germline mutations in fundamental epigenetic regulatory molecules including DNA methyltransferase 3A (*DNMT3A*) are commonly associated with growth disorders, whereas somatic mutations are often associated with malignancy. We profiled genome-wide DNA methylation patterns in *DNMT3A* c.2312G>A; p.(Arg771Gln) carriers in a large Amish sibship with Tatton-Brown-Rahman syndrome (TBRS), their mosaic father and 15 TBRS patients with distinct pathogenic de novo *DNMT3A* variants. This defined widespread DNA hypomethylation at specific genomic sites enriched at locations annotated to genes involved in morphogenesis, development, differentiation, and malignancy predisposition pathways. TBRS patients also displayed highly accelerated DNA methylation aging. Notably, these findings were most striking in a carrier of the AML associated driver mutation p.Arg882Cys. Our studies additionally defined phenotype related accelerated and decelerated epigenetic aging in two histone methyltransferase disorders; NSD1 Sotos syndrome overgrowth disorder and KMT2D Kabuki syndrome growth impairment. Together, our findings provide fundamentally new insights into aberrant epigenetic mechanisms, the role of epigenetic machinery maintenance and determinants of biological aging in these growth disorders.

## Introduction

DNA methylation is an essential epigenetic process involving the addition of a methyl group to cytosine. It is known to play a role in many important genomic regulatory processes, including X-chromosome inactivation, genomic imprinting and the repression of tumor suppressor genes in cancer, mediating transcriptional regulation as well as genomic stability (Jones 2012). Three catalytically active DNA methyltransferases (DNMTs) are involved in the methylation of cytosine: *DNMT1*, which is mainly responsible for the maintenance of DNA methylation over replication and *DNMT3A* and *DNMT3B*, which generally perform *de novo* methylation of either unmethylated or hemimethylated DNA. An absence of these enzymes in mice results in embryonic (*DNMT1* and *3B*) or postnatal (*DNMT3A*) lethality (Okano et al. 1999), confirming their essential roles in development. In line with knockout mouse models, pathogenic variants affecting the chromatin binding domains of *DNMT1* have been shown to cause two separate progressive autosomal dominant adult-onset neurologic disorders (Klein et al. 2011). Biallelic pathogenic variants in *DNMT3B* have been associated with immunodeficiency, centromere instability and facial anomalies (ICF) syndrome (Jiang et al. 2005). To date, *DNMT3A* has been linked to a number of physiological functions, including cellular differentiation, malignant disease, cardiac disease, learning and memory formation. Somatically acquired pathogenic variants in *DNMT3A* are associated with over 20% of acute myeloid leukemia (AML) cases, whilst heterozygous germline pathogenic variants have more recently been found to underlie Tatton-Brown-Rahman syndrome (TBRS; also known as DNMT3A-overgrowth syndrome, OMIM: 615879) (Challen et al. 2011; Tatton-Brown et al. 2014). TBRS is characterized by increased growth, intellectual disability (ID) and dysmorphic facial features.

There is an emerging group of epigenetic regulatory molecule-associated human growth disorders where the underlying molecular defect is a disruption to the DNA methylation and histone machinery. There are now over 40 disorders identified within this group, which can be further subgrouped into diseases resulting from disruption of the ‘writers’, ‘readers’ and ‘erasers’ of epigenetic modifications (Bjornsson 2015). Example disorders in each group include Kabuki, Sotos and Weaver syndromes (‘writers’), Smith-Magenis, Rett and Bohring-Optiz syndromes (‘readers’), and Wilson-Turner and Cleas-Hensen syndromes (‘erasers’). The final subgroup occurs due to disruption of chromatin remodellers, with example resulting disorders including CHARGE and Floating Harbour syndromes. Neurological and cognitive impairment are common features of these conditions, suggesting that precise epigenetic regulation may be critical for neuronal homeostasis. However, a true understanding of the pathogenic mechanism underlying these conditions remains poorly understood.

In the current study, we investigated the methylomic consequences of a *DNMT3A* pathogenic variant (NC_000002.12:g.25240312C>T; NM_022552.4:c.2312G>A; p.Arg771Gln) in a large Amish family comprising four individuals affected with TBRS arising due to a mosaic pathogenic *DNMT3A* variant in their father (Xin et al. 2017). The occurrence of multiple affected and unaffected individuals in the same sibship, together with the combined genetic and environmental homogeneity of the Amish, permitted an in-depth investigation of the genome-wide patterns of DNA methylation associated with pathogenic variation in *DNMT3A*. We subsequently extended our analyses to other (non-Amish) TBRS patients harbouring distinct pathogenic *de novo DNMT3A* variants, as well other methyltransferase associated overgrowth and growth deficiency syndromes, defining altered epigenetic profiles as common key themes of these growth disorders.

## Results

### Reduced DNA methylation at sites involved in morphogenesis, development and differentiation in TBRS patients

*DNMT3A* encodes a DNA methyltransferase with both *de novo* and maintenance activity (Okano et al. 1999; Chen et al. 2003). We first looked for global changes in DNA methylation in whole blood obtained from *DNMT3A* c.2312G>A; p.(Arg771Gln) carriers, using the methylation-sensitive restriction enzyme based LUMA assay (Karimi et al. 2006) to quantify DNA methylation across GC rich regions of the genome, finding no evidence for altered global DNA methylation (LUMA: mean *DNMT3A* c.2312G>A carriers = 0.274, wildtype = 0.256, t-test p-value = 0.728). We next quantified DNA methylation at 414,172 autosomal sites across the genome using the Illumina 450k array. Globally, no difference in mean DNA methylation was found in any of the *DNMT3A* heterozygous c.2312G>A; p.(Arg771Gln) samples tested (Wilcoxon Rank Sum test p-value < 2.2×10^−16^). In contrast, an analysis of site-specific DNA methylation differences in *DNMT3A* c.2312G>A; p.(Arg771Gln) heterozygotes (including the mosaic father) vs wildtype individuals in the Amish pedigree identified 2,606 differentially methylated positions (DMPs) (Benjamini-Hochberg FDR < 0.05) (**Figure 1a,b** and **Supplemental Table S1, Supplemental Figure S1**), of which 1,776 DMPs were characterized by a >10% change in DNA methylation. Interestingly, these DMPs were highly enriched for sites characterized by reduced DNA methylation in *DNMT3A* c.2312G>A; p.(Arg771Gln) heterozygotes (n = 2,576 DMPs, 98.85%, sign-test p-value < 2.2×10^−16^). We examined the extent to which these findings were potentially driven by cell-type differences between *DNMT3A* c.2312G>A; p.(Arg771Gln) carriers and wildtype family members using derived blood cell-type estimates from each sample (Horvath 2013). There was a highly significant correlation (r = 0.876, p-value < 2.2×10^−16^, **Supplemental Figure S2**) in effect sizes at the 2,606 DMPs between models including and excluding cell-types as covariates, indicating that the observed patterns of differential DNA methylation are not strongly influenced by cell-type variation. We used DMRcate (Peters et al. 2015) to identify spatially correlated regions of differential DNA methylation significantly associated with the *DNMT3A* c.2312G>A; p.(Arg771Gln) variant, identifying 388 autosomal differentially methylated regions (DMRs) (example DMR shown in **Supplemental Figure S3**), all characterized by hypomethylation in *DNMT3A* c.2312G>A; p.(Arg771Gln) carriers apart from one 739bp DMR which showed increased DNA methylation (**Supplemental Table S2**). The mean size of the identified DMRs was 625bp (range = 6 to 5,522bp), spanning an average of 6 probes (**Supplemental Figure S4**).

**Figure 1.**
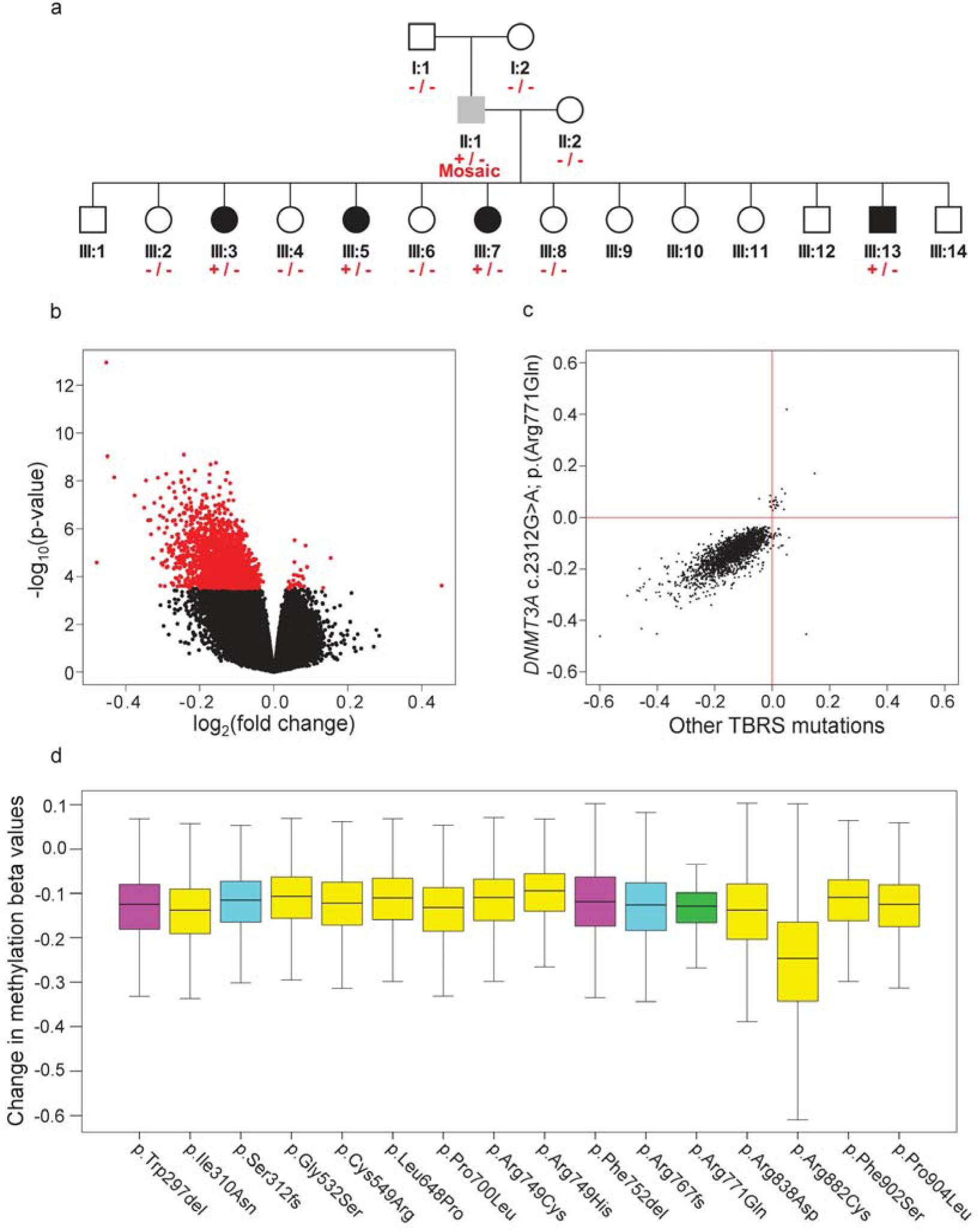
TBRS *DNMT3A* variants are associated with widespread DNA hypomethylation. (a) Simplified pedigree indicating the genotyping of individuals in the Amish family investigated (‘+/-‘; carriers of the *DNMT3A* c.2312G>A p.[Arg771Gln] variant, ‘+/- Mosaic’; the father, and ‘-/-‘; wildtype individuals). Black shading; individuals with a phenotype consistent with TBRS, grey shading; the father with macrocephaly and mild intellectual impairment, and white shading; unaffected individuals. Each of these samples was profiled on the Illumina 450k DNA methylation array. (b) Volcano plot showing site-specific DNA methylation differences (x-axis) and –log10 P values (y-axis) from an analysis comparing Amish *DNMT3A* c.2312G>A; p.(Arg771Gln) pathogenic variant carriers and wildtype family members using the Illumina 450K array. Red values indicate the 2,606 differentially methylated positions (DMPs) detected at a Benjamini Hochberg FDR < 0.05. (c) Comparison of *DNMT3A* c.2312G>A; p.(Arg771Gln) identified differentially methylated positions (log_2_ fold change) relative to other *DNMT3A* TBRS associated variants assessed in this study (all variants grouped and measured relative to controls). Pearson correlation = 0.7642, p-value < 2.2 x 10^−16^. (d) Boxplot illustrating the DNA methylation changes observed in association with the *DNMT3A* TBRS variants studied at the differentially methylated positions identified in the Amish *DNMT3A* c.2312G>A p.[Arg771Gln] carriers. The predicted protein consequence of each *DNMT3A* variant studied is indicated (pink; in-frame deletion, yellow; single nucleotide variant, cyan; duplications predicted to result in a frameshift, green; Amish c.2312G>A; p.(Arg771Gln) variant).

We next investigated whether DNMT3A p.(Arg771Gln)-associated DMPs are enriched in specific genic locations (see **Methods**). We found a modest enrichment of DMPs in regions 1500bp or more upstream of the transcriptional start site (Chi-Squared Yates corrected p-value=0.047) and more prominent enrichment in intergenic regions (Chi-Squared Yates corrected p-value=1.54×10^−14^) (**Supplemental Figure S5A**). DMPs were also significantly enriched in CpG Island shore regions (Chi-Squared Yates corrected p-value=8×10^−30^) (**Supplemental Figure S5B**). We also examined DMP occurrence in experimentally determined cancer and reprogramming specific DMR locations (Doi et al. 2009), finding a 2.4 fold and 4.7 fold overrepresentation (Chi-Squared Yates corrected p-value=4.24×10^−6^ and Chi-Squared Yates corrected p-value<1.3×10^−41^ respectively), as well as predicted enhancer elements which showed a 1.4 fold overrepresentation (Chi-Squared Yates corrected p-value=7.57×10^−16^). We then undertook gene ontology analysis, accounting for the background distribution of probes on the Illumina 450K array, to functionally annotate the DNA methylation differences observed in the carriers. The 2,606 DMPs identified in this study showed significant overrepresentation in functional pathways related to morphogenesis, development and differentiation (top hit: GO:0007275 -multicellular organism development, contains 474 genes associated with DMPs, FDR Q-value=3.7×10^−7^) (**Supplemental Table S3**).

To provide additional evidence to support the notion that *DNMT3A* c.2312G>A; p.(Arg771Gln) carriers exhibit disruption to developmental pathways, we used the Genomic Regions Enrichment of Annotation Tool (GREAT) (McLean et al. 2010) to explore functional pathways enriched in genes annotated to *DNMT3A* c.2312G>A; p.(Arg771Gln)-associated DMRs. This revealed a significant effect on genes implicated in developmental pathways (1^st^ ranked GO Biological Process = Skeletal System Development, fold enrichment = 2.5, binomial FDR Q-value=9.22×10^−7^), with a specific enrichment for Homeobox protein domain encoding genes (InterPro) (fold enrichment=239.59, binomial FDR Q-value=4.15×10^−23^), fundamental for normal developmental processes as well as a number of malignancy terms (from the Molecular Signatures Database (Liberzon et al. 2011)) (1^st^ ranked term = Genes with promoters occupied by PML-RARA fusion protein in acute promyelocytic leukemia (APL) cells NB4 and two APL primary blasts, based on Chip-seq data, fold enrichment=3.14, binomial FDR Q-value=6.18×10^−7^) (see **Supplemental File S1**).

To establish whether these differentially methylated positions are a consistent feature of TBRS, we profiled a further 15 non-Amish patients carrying distinct previously published *DNMT3A* pathogenic variants (see **Figure 2** and **Table 1**) using the Illumina EPIC DNA methylation array. Importantly, examination of the originally differentially methylated positions identified in *DNMT3A* c.2312G>A; p.(Arg771Gln) mutation carriers revealed that the majority of differentially methylated positions were common to all of the TBRS patients regardless of the underlying causative *DNMT3A* variant (**Figure 1c**), with a Pearson correlation coefficient of 0.7642 (p-value<2.2×10^−16^) for effect sizes across all DMPs. Each variant showed some heterogeneity in effect size (**Figure 1d**), with *DNMT3A* c.2644C>T p.(Arg882Cys) associated with the greatest overall changes in DNA methylation. These data lead us to conclude that TBRS patients show loss of methylation at sites annotated to genes involved in development and growth pathways, mirroring the well-characterized overgrowth and neurocognitive features that characterize this disorder.

**Table 1.** *DNMT3A* genotype (p.Arg771Gln shown in blue; p.Arg882Cys, shown in red), chronological age, predicted epigenetic age and percentage of age acceleration calculated for TBRS syndrome cases included in this study, * Age acceleration taken from linear regression model applied to 4 individuals carrying the mutation.

| ID | Mutation<br>(nucleotide) | Mutation<br>(protein) | Chronological<br>Age (years) | Epigenetic<br>Age<br>(years) | Epigenetic<br>Age<br>Acceleration<br>(fold<br>change) | Epigenetic Age<br>Acceleration<br>(percentage<br>increase) |
| --- | --- | --- | --- | --- | --- | --- |
| **Single nucleotide variants** |  |  |  |  |  |  |
| 1 | c.929T>A | p.Ile310Asn | 9.27 | 24.2 | 2.61 | 161% |
| 2 | c.1594G>A | p.Gly532Ser | 5.92 | 22.7 | 3.83 | 283% |
| 3 | c.1645T>C | p.Cys549Arg | 9.36 | 25 | 2.67 | 167% |
| 4 | c.1943T>C | p.Leu648Pro | 19.34 | 32.1 | 1.66 | 66% |
| 5 | c.2099C>T | p.Pro700Leu | 13.45 | 32.5 | 2.42 | 142% |
| 6 | c.2245C>T | p.Arg749Cys | 13.87 | 19.6 | 1.41 | 41% |
| 7 | c.2246G>A | p.Arg749His | 8.9 | 33.6 | 3.78 | 278% |
| 8 | **c.2312G>A** | **p.Arg771Gln** | **8 - 23** | **15.8 - 36.1** | **1.41** | **41%\*** |
| 9 | c.2512A>G | p.Asn838Asp | 14.44 | 38.8 | 2.69 | 169% |
| 10 | **c.2644C>T** | **p.Arg882Cys** | **2.28** | **21.9** | **9.61** | **861%** |
| 11 | c.2705T>C | p.Phe902Ser | 9.84 | 25.4 | 2.58 | 158% |
| 12 | c.2711C>T | p.Pro904Leu | 7.78 | 35.7 | 4.59 | 359% |
| **In-frame deletions** |  |  |  |  |  |  |
| 13 | c.889_891delTGG | p. Trp297del | 5.82 | 23.2 | 3.99 | 299% |
| 14 | c.2255_2257delTCT | p.Phe752del | 3.05 | 10.6 | 3.48 | 248% |
| **Duplications resulting in a frameshift** |  |  |  |  |  |  |
| 15 | c.2297dupA | p.Arg767fs | 3.12 | 14.4 | 4.62 | 362% |
| 16 | c.934_937dupTCTT | p.Ser312fs | 21 | 38.9 | 1.85 | 85% |

**Figure 2.**
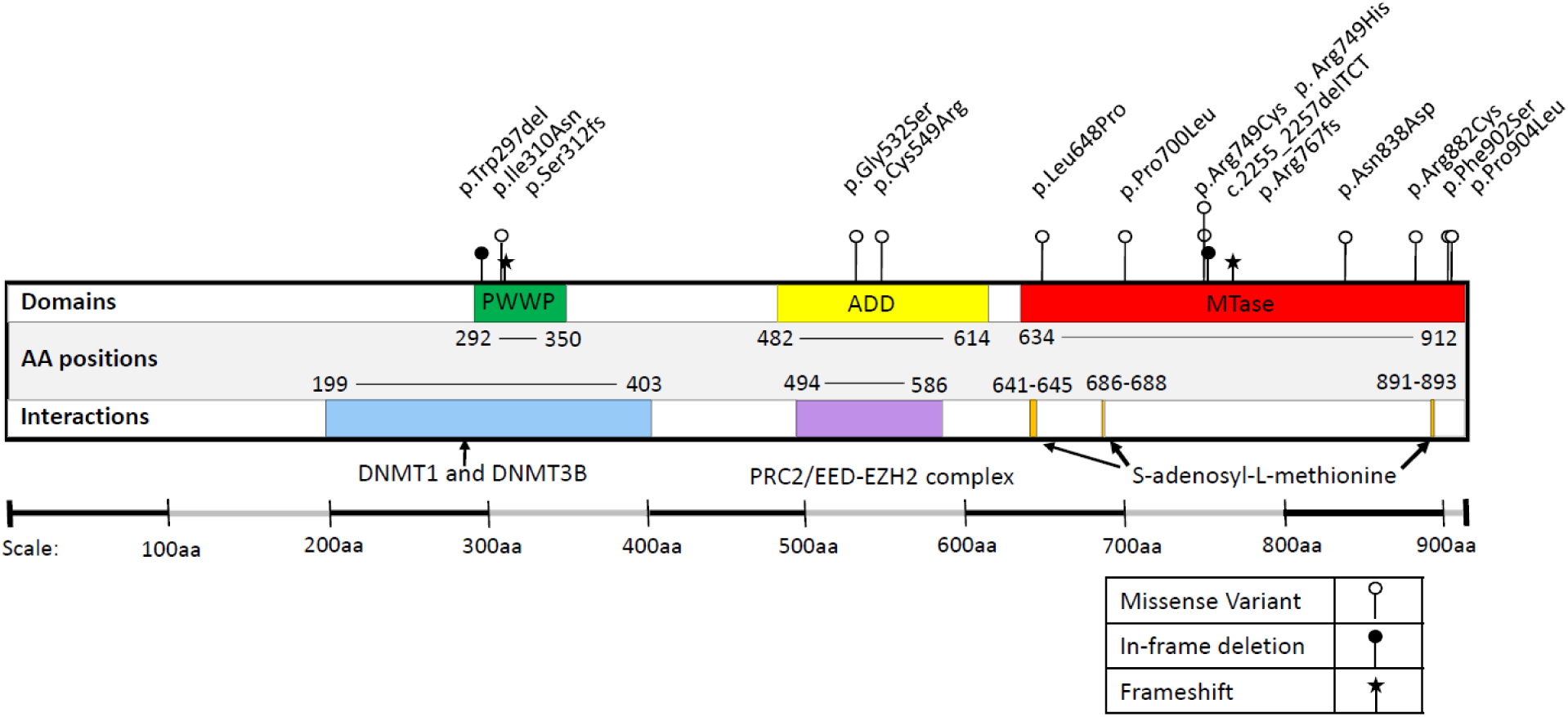
Schematic representation of DNMT3A. The positions of the disease-associated variants included in this study are indicated relative to the protein domain architecture.

### DNMT3A mutations are associated with highly accelerated epigenetic aging, particularly the cardinal AML driver mutation p.Arg882Cys

DNA methylation at a specific set of CpG sites, representing a so-called “epigenetic clock”, has been shown to be strongly correlated with chronological age (Horvath 2013). Deviations from chronological age have been associated with several measures of accelerated biological aging and age-related phenotypes (Johnson et al. 2012; Levine et al. 2015; Marioni et al. 2015; Chen et al. 2016). We investigated the DNA methylation age of *DNMT3A* c.2312G>A p.(Arg771Gln) carriers using the DNA age calculator (http://dnamage.genetics.ucla.edu/) (Horvath 2013) finding that *DNMT3A* c.2312G>A; p.(Arg771Gln) carriers show evidence for highly-accelerated aging - an increase of ~40% beyond their chronological age - when compared to wildtype family members (ANCOVA p-value=0.004) (**Figure 3a**). Consistent with this, the mosaic father was found to have intermediate level of epigenetic age acceleration, with a 23% increase over his chronological age. This age acceleration was a cumulative process as indicated by the increased slope of *DNMT3A* c.2312G>A; p.(Arg771Gln) carriers vs wildtype. Epigenetic age could therefore be predicted by the linear regression model as follows: epigenetic age = 4.81 + 1.405 x chronological age. The cumulative increase of epigenetic age relative to chronological age is also notable when compared to a recent meta-analysis of longitudinal cohort data which shows the trajectory of epigenetic age in different populations progresses at a slightly slower rate when compared to increasing chronological age (Marioni et al. 2018).

**Figure 3.**
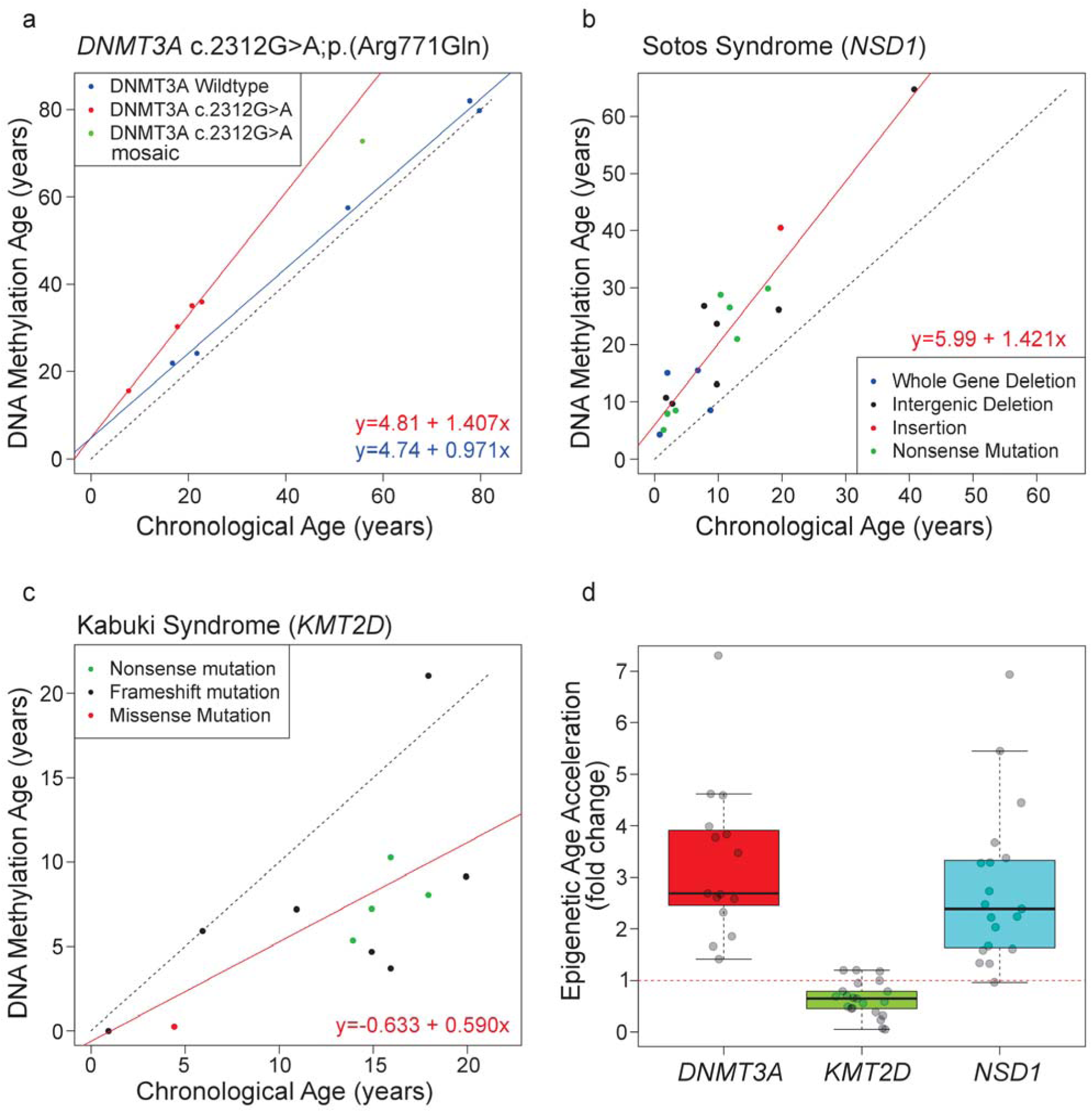
Altered epigenetic aging is observed in methyltransferase-associated human growth disorders. (a) Scatter plot comparing ‘DNA methylation age’ derived from the Illumina 450K data (y-axis) and actual chronological age (x-axis) in *DNMT3A* c.2312G>A p.(Arg771Gln) pathogenic variant carriers (red) vs wildtype family members (blue). Green indicates the mosaic individual. The linear regression model is also shown. (b) Scatter plot comparing DNA methylation age vs chronological age in patients with Sotos syndrome. Inframe legend illustrates the different *NSD1* pathogenic variant studied. (c) Scatter plot comparing DNA methylation age vs chronological age in patients with Kabuki syndrome. Inframe legend illustrates the different *KMT2B* pathogenic variant studied. (d) Boxplot comparing the epigenetic age acceleration rates found in association with TBRS *DNMT3A* variants, *KMT2D* Kabuki syndrome variants and *NSD1* Sotos syndrome variants. Each age acceleration observation is plotted as a circle. The dotted red line denotes no age acceleration.

We next tested for evidence of elevated epigenetic aging in the 15 additional *de novo DNMT3A* pathogenic variant carriers with TBRS overgrowth syndrome. All patients showed accelerated epigenetic aging, although the position and type of each variant results in differing degrees of accelerated epigenetic aging (**Table 1**). Notably, the greatest rate of epigenetic age acceleration (>800%) was observed in association with the germline p.(Arg882Cys) substitution, somatic mutation of DNMT3A Arg882 being the most commonly associated with acute myeloid leukemia (AML).

### Altered epigenetic aging in methyltransferase-associated human growth disorders

To determine whether altered epigenetic aging is a characteristic of other growth disorders associated with disruption of epigenetic regulatory molecules, we extended our study using publicly-available Illumina 450K DNA methylation data. We first analysed the data from patients with Sotos syndrome, a congenital overgrowth syndrome which results from mutation of the epigenetic modifier *NSD1* (**Supplemental Table S4**), a lysine histone methyltransferase, (Kurotaki et al. 2002; Qiao et al. 2011). Notably, consistent with *DNMT3A* pathogenic variant carriers, these individuals are characterized by an epigenetic age acceleration of ~40% (linear regression model R^2^=0.869, p-value=6.4×10^−9^) (see **Figure 3b**). We then examined data from patients with loss of function variants in the *KMT2D* gene (**Supplemental Table S5**), also encoding a lysine histone methyltransferase, which causes Kabuki syndrome (Ng et al. 2010; Butcher et al. 2017), a multisystem disorder with patients typically presenting with post-natal growth deficiency (rather than overgrowth), characteristic facial features, intellectual disability and other variable features. Although there is more heterogeneity in epigenetic age when compared with the *NSD1* mutation carriers there was a significant reduction in epigenetic age of approximately 40% seen across these patients (linear regression model R^2^=0.418, p-value=0.023) (**Figure 3c)**.

## Discussion

To date, 78 individuals have been described with the overgrowth condition TBRS. Within this group a wide variety of germline *DNMT3A* pathogenic variants have been reported including 33 missense, eight stop-gain, seven frameshift, two splice site variants, two in-frame and five whole-gene deletions (including a set of identical twins) (Tatton-Brown et al. 2014; Okamoto et al. 2016; Tlemsani et al. 2016; Hollink et al. 2017; Kosaki et al. 2017; Lemire et al. 2017; Shen et al. 2017; Spencer et al. 2017; Tatton-Brown et al. 2017; Xin et al. 2017; Tatton-Brown et al. 2018). Clinically, the predominant features of TBRS are overgrowth, a characteristic facial gestalt and neurocognitive impairment. These features show phenotypic overlap with conditions associated with germline pathogenic variants in other epigenetic regulatory genes, including Sotos and Weaver syndrome caused by variants in *NSD1* and *EZH2* histone methyltransferases respectively (Tatton-Brown et al. 2017). These genes encode essential epigenetic regulatory proteins, with a dual somatic/germline role in the pathogenesis of haematological malignancies and overgrowth syndromes with variable degrees of intellectual impairment (Tatton-Brown et al. 2014).

The majority of *DNMT3A* pathogenic variants in TBRS have been found to be *de novo*, with five individuals inheriting the pathogenic variant from two mosaic parents (Tlemsani et al. 2016; Xin et al. 2017) and two individuals inheriting the pathogenic variant from their affected father (Lemire et al. 2017). Extensive studies of the role of *DNMT3A* in haematopoietic stem cell (HSC) differentiation are also reported, including the regular occurrence of somatic *DNMT3A* variants in patients with acute myeloid leukaemia (AML). The most common somatic pathogenic variant reported in patients with AML affects the amino acid residue Arg882. To date, pathogenic variation at this residue has been described in the germline of 12 TBRS patients, five with p.(Arg882His) and seven with p.(Arg882Cys) (Tlemsani et al. 2016; Hollink et al. 2017; Kosaki et al. 2017; Shen et al. 2017; Spencer et al. 2017; Tatton-Brown et al. 2018). Despite these studies, the underlying biological mechanism and outcomes of gene mutation in TBRS, and the potential risks of haematological malignancy, remain largely unclear.

Here we investigated variation in DNA methylation associated with a germline heterozygous *DNMT3A* missense pathogenic variant c.2312G>A; p.(Arg771Gln), affecting the catalytic MTase domain, in a large Amish family comprising four children with TBRS, unaffected siblings, and their mosaic father who displayed an intermediate clinical phenotype (Xin et al. 2017). Affected individuals were characterized by widespread hypomethylation, with DMPs enriched in the vicinity of genes associated with growth and development, tissue morphogenesis and differentiation. These differences were also observed in an analysis of 15 patients carrying differing *de novo* pathogenic variants in *DNMT3A*. While dysregulation of growth control has been linked to numerous developmental disorders and malignancy, the specific molecular basis of this relationship is not fully understood. Our finding of altered epigenetic outcomes in TBRS prompted us to consider similar investigations in other growth disorders associated with epigenetic dysfunction; Sotos syndrome, a neurodevelopmental disorder with features overlapping TBRS and associated with overgrowth in childhood due to histone methyltransferase *NSD1* gene alterations, and Kabuki syndrome, a distinct neurodevelopmental disorder associated with poor growth and histone methyltransferase *KMT2D* gene alterations. Remarkably we defined clear aberrations in epigenetic aging appropriate to the specific nature of each condition. In both overgrowth conditions, TBRS and Sotos syndrome, we identified accelerated epigenetic aging as measured by the DNA methylation age calculator (Horvath 2013). Conversely patients with Kabuki syndrome clinically characterized by poor growth, displayed decelerated epigenetic age. Epigenetic age has been strongly correlated with chronological age in unaffected individuals in previous studies of a variety of tissue types (Hannum et al. 2013; Horvath 2013). The observation of accelerated epigenetic aging in both TBRS and Sotos syndrome potentially results from reduced methyltransferase activity, in addition to increased cell turnover associated with the overgrowth seen with these disorders, with the converse being the case for Kabuki syndrome. Accelerated epigenetic aging has been associated with age-related clinical characteristics and mortality in epidemiological studies. For example, accelerated epigenetic age in lymphocytes correlates with reduced physical and cognitive function in the elderly, and increased overall mortality independent of other variables such as BMI, sex and smoking status (Marioni et al. 2015; Chen et al. 2016). Accelerated epigenetic age has also been found in specific diseases such as Huntington’s disease (+ 3.4 years) (Horvath et al. 2016), Down’s syndrome (+ 6.6 years) (Horvath et al. 2015) and Werner’s syndrome (+ 6.4 years) (Maierhofer et al. 2017). Remarkably, unlike these other disorders, a distinguishing feature of carriers of the Amish *DNMT3A* c.2312G>A; p.(Arg771Gln) variant is that comparison of epigenetic age vs chronological age across the age ranges available is supportive of a marked cumulative acceleration of epigenetic age over life time course. While these studies were not possible for the other *DNMT3A* variants, this may be indicative of a similar effect on cumulative epigenetic age acceleration over life course in TBRS.

There are only four reported cases of an AML tumour carrying the DNMT3A p.Arg771Gln mutation which was present as a germline mutation in the Amish TBRS patients in our study. Biochemical measurements of DNMT3A enzyme show that mutations at both the Arg771 and Arg882 residues result in reduced methyltransferase activity, with a greater degree of reduction resulting from Arg882 variants compared to Arg771 variants (2.4 fold difference) (Holz-Schietinger et al. 2012). Given this reduced methyltransferase activity, we may expect to observe more pronounced changes in DNA methylation in patients with germline variants affecting Arg882 when compared to variants affecting other amino acid residues such as Arg771. Our data reflected this notion, with alteration Arg882 displaying markedly greater methylation changes compared to the other DNMT3A mutations investigated in this study. Currently available literature suggests that the risk of haematological malignancy in TBRS individuals may vary depending on the specific pathogenic variant underling their condition (Hollink et al. 2017). The significantly advanced epigenetic age that we observed in association with p.Arg882Cys may explain why haematological malignancy has to date only been reported in two TBRS patients, one harbouring this germline variant and the second the p.Tye735Ser variant, the latter not being assessed in this study (Hollink et al. 2017; Tatton-Brown et al. 2018).

In summary, our findings identify widespread DNA hypomethylation in genes involved in morphogenesis, development, differentiation and malignancy in TBRS patients. These findings provide important new insights into the role of DNMT3A during development, and of relevance to haematological malignancy, and define perturbation to epigenetic machinery and biological ageing as common themes in overgrowth and growth deficiency syndromes.

## Methods

### Genetic and Clinical Studies

The phenotypic features of the four affected siblings (3 females and 1 male, aged 11-25 years; Figure 1a individuals III:3, III:5, III:7 and III:13) include: macrocephaly, tall stature, obesity, hypotonia, mild to moderate intellectual disability, behavioural problems, and a distinctive facial appearance. Whole genome SNP genotyping and exome sequencing of DNA samples taken with informed consent under regionally-approved protocols excluded mutations in known genes, or candidate new genes, associated with neurodevelopmental disorders. Subsequent studies defined a heterozygous c.2312G>A substitution in *DNMT3A,* resulting in a p.(Arg771Gln) substitution associated with TBRS as the cause of the condition, and full clinical details are described in Xin et al 2017 (Xin et al. 2017). Further testing revealed mosaicism for the *DNMT3A* c.2312G>A variant in the father, and Xin et al. (Xin et al. 2017), demonstrated pathogenic variant load varied in different tissue types.

### DNA Methylation Profiling

Genomic DNA from blood was sodium bisulfite converted using the EZ-96 DNA Methylation Kit (Zymo Research, Irvine, CA, USA) and DNA methylation quantified across the genome using the Illumina Infinium HumanMethylation450 array (“Illumina 450K array”) (Illumina, San Diego, CA, USA). The additional 15 *DNMT3A* pathogenic variants were profiled using the Illumina Infinium EPIC array (Illumina, San Diego, CA, USA). The Bioconductor package *wateRmelon* (Pidsley et al. 2013) in R 3.4.1 was used to import idat files and, after checking for suitable signal intensities and sodium bisulfite conversion, the DNA methylation data was quantile normalized using the *dasen* function in *wateRmelon* and methylation beta values produced (ratio of intensities for methylated vs unmethylated alleles). Probes showing a detection p-value >0.01 in at least 1% of samples or a beadcount <3 in 5% of samples were removed across all samples. Probes containing common SNPs within 10 bp of the CpG site were removed (minor allele frequency > 5%). Nonspecific probes and probes on the sex chromosomes were also removed (Chen et al. 2013; Price et al. 2013). Differentially methylated positions (DMPs) were identified using a *limma* based linear model based on pathogenic variant genotype and sex as a covariate (Smyth 2004). Significance was determined using the Benjamini-Hochberg false discovery rate (FDR) of 5% (Benjamini and Hochberg 1995). Changes in methylation were calculated based on comparison between *DNMT3A* c.2312G>A; p.(Arg771Gln) carriers vs wildtype individuals in the Amish pedigree. The additional 15 *DNMT3A* pathogenic variants were assessed relative to 7 anonymous control samples run on the same EPIC array run. Blood cell counts were unknown and so were estimated using the DNA methylation age calculator (Horvath 2013; Koestler et al. 2013), and assessed in the linear model. To identify differentially methylated regions (DMRs), the package DMRcate was used with the same *limma* based design (Peters et al. 2015). Gene ontology enrichment analysis was performed using genes annotated to FDR corrected differentially methylated positions using the *gometh* function of the *missMethyl* package (Phipson et al. 2016) which takes into account potential bias of probe distributions on the beadchip array. KEGG pathway analysis was performed using the *gsameth* command of *missMethyl* and KEGG annotation files from the Bioconductor KEGGREST package (http://bioconductor.org/packages/release/bioc/html/KEGGREST.html). Regional enrichment analysis based on Illumina annotations was performed using a Chi-squared test with Yates correction in R. Differentially methylated regions were functionally annotated using the webtool GREAT (http://great.stanford.edu/public/html/).

Global methylation measurements were made using the Luminometric methylation assay (LUMA) (Karimi et al. 2006) based on cleavage by a methylation sensitive restriction enzyme followed by polymerase extension assay via pyrosequencing on the Pyromark 24 (Qiagen). Epigenetic age calculations were made using the DNA methylation age calculator (https://dnamage.genetics.ucla.edu/) for Illumina 450k data, and Illumina EPIC arrays assessed using the agep function of the *wateRmelon* Bioconductor package, the latter based on the original calculator developed by Steve Horvath (Horvath 2013). Accelerated age was calculated for the Amish TBRS *DNMT3A* c.2312G>A; p.(Arg771Gln), Sotos syndrome and Kabuki syndrome patients based on linear models of recorded chronological age and calculated epigenetic age. Estimates of age acceleration for the additional 15 TBRS cases were calculated by dividing the calculated epigenetic age with their chronological age. Additional Sotos syndrome patient Illumina DNA methylation files were obtained from GEO accession GSE74432, with corresponding chronological ages derived from the associated paper (Choufani et al. 2015). Kabuki syndrome DNA methylation data and chronological age was obtained from GEO accession GSE97362 (Butcher et al. 2017).

## Supporting information

## Acknowledgements

We are grateful to the Amish families for participating in this study, and to the Amish community for their continued support of the Windows of Hope project. The work was supported by the Newlife Foundation for Disabled Children (Ref: SG/16-17/02, to C.G.S, A.R.J, A.H.C and E.L.B), MRC grant G1001931 (to E.L. Baple), MRC grant G1002279 (to A.H. Crosby) and MRC grants MR/M008924/1 and MR/K013807/1 (to J. Mill).

## Contributions

E.L.B. A.H.C and J.M. conceived study. R.M. and B.A.C. performed the genetic analysis. A.R.J. and R.M. performed LUMA global methylation assay. A.R.J. performed the microarray based DNA methylation analysis and all associated statistical analyses. A.R.J., C.G.S., R.M., A.H.C.,

J.M. and E.L.B. wrote the manuscript, H.E.C. and M.A.P., D.M., F.I.R., K.T.B., L.A., D.B., T.D., M.F.H., R.N.E., A.K., A.K., A.S., A.S., L.V.M., V.V., and N.Y. contributed samples and clinical data.

## Disclosure Declaration

The authors declare that there are no conflicts of interest.

