## Supplementary material for "Growth disrupting mutations in epigenetic regulatory molecules are associated with abnormalities of epigenetic aging"

**Whole gene deletions**

| Sample ID | Exon | Mutation | Protein change | Inheritance | Sex | Age at sample collection (years) |
|---|---|---|---|---|---|---|
| 11D_0326 | 2Mb deletion | chr5:175,366,008-177,470,488 (hg19) | | *De novo* | F | 9 |
| 11D_0328 | 1.3 Mb deletion | chr5:175,764,262-177,059,256 (hg19) | | *De novo* | F | 7 |
| DL151889 | 2Mb deletion | microdeletion of distal 5q35.2 | | *De novo* | M | 2.2 |
| DL87406 | 1.9Mb deletion | 5q35.2-35.3 (RP11-67P18 to RP11-423H2), FISH BAC RP11-99N22 deleted | | *De novo* | M | <1 |

**Intragenic deletions**

| Sample ID | Exon | Mutation | Protein change | Inheritance | Sex | Age at sample collection (years) |
|---|---|---|---|---|---|---|
| DL38402 | 5 | c.1583delA | p.Lys528Argfs*8 | *De novo* | M | 19.7 |
| DL50448 | 5 | c.2014-2018delACAGA | p.Thr672Glufs*9 | Inherited from father | M | 8 |
| DL50450 | 5 | c.2014-2018delACAGA | p.Thr672Glufs*9 | *De novo* | M | 41 |
| DL50452 | 5 | c.2014-2018delACAGA | p.Thr672Glufs*9 | Inherited from father | F | 2 |
| 11D/6718 | 5 | c.1716delC | p.Cys573Valfs*26 | *De novo* | F | 10 |
| DL122057 | 13 | c.4843delT | p.Tyr1615Thrfs*27 | *De novo* | M | 3 |
| 11D/6637 | 15-19 | ex15-19 del | | *De novo* | M | 10 |

**Insertion**

| Sample ID | Exon | Mutation | Protein change | Inheritance | Sex | Age at sample collection (years) |
|---|---|---|---|---|---|---|
| DL94609 | 14 | 4977_4978insG | p.Arg1660Alafs*13 | *De novo* | M | 20 |

**Nonsense mutations**

| Sample ID | Exon | Mutation | Protein change | Inheritance | Sex | Age at sample collection (years) |
|---|---|---|---|---|---|---|
| DL159249 | 22 | c.6349C>T | p.Arg2117* | *De novo* | F | 12 |
| DL168744 | 5 | c.1492C>T | p.Arg498* | *De novo* | M | 2.2 |
| DL89813 | 5 | c.1801A>T | p.Lys601* | *De novo* | M | 10.6 |
| DL117330 | 16 | c.5445C>G | p.Tyr1815* | *De novo* | F | 13.2 |
| DL76010* | 5 | c.1810C>T | p.Arg604* | | F | 1.6 |
| DL179067 | 22 | c.6454C>T | p.Arg2152* | *De novo* | M | 18 |
| A1208 | 22 | c.6454C>T | p.Arg2152* | *De novo* | F | 3.5 |

**Supplemental Table 4** – Sotos syndrome patients with pathogenic *NSD1* loss of function variants (taken from Supplemental Data 1 of Choufani et al. Nature Communications 2015, doi:10.1038/ncomms10207)
