## Supplementary material for "Growth disrupting mutations in epigenetic regulatory molecules are associated with abnormalities of epigenetic aging"

***KMT2D* pathogenic loss of function variants**

| Sample ID | mutation DNA | mutation protein | Coding effect |
|---|---|---|---|
| KMT2D-1 | c.15061C>T | p.Arg5021* | nonsense |
| KMT2D-2 | c.16318delG | p.Glu5440Argfs*16 | frameshift |
| KMT2D-3 | c15030dupA | p.Glu5011Argfs*13 | frameshift |
| KMT2D-4 | c.8172_8173delC | p.Phe2724Glnfs*5 | frameshift |
| KMT2D-5 | c.6595delT | p.Tyr2199Ilefs*65 | frameshift |
| KMT2D-6 | c.14055-14056delCA | p.His4685Glnfs*4 | frameshift |
| KMT2D-7 | c.6295C>T | p.Arg2099* | nonsense |
| KMT2D-8 | c.4135_4136delA | p.Met1379Valfs*52 | frameshift |
| KMT2D-9 | c.12592C>T | p.Arg4198* | nonsense |
| KMT2D-10 | c.4135_4136delA | p.Met1379Valfs*52 | frameshift |
| KMT2D-11 | c.11710C>T | p.Gln3904* | nonsense |

***KMT2D* missense variant**

| Sample ID | mutation DNA | mutation protein | Coding effect/Splice site mutation |
|---|---|---|---|
| KMT2D-12 | c.15143G>A | p.Arg5048His | missense |

**Supplemental Table 5** – Kabuki syndrome patients with pathogenic *KMT2D* variants (Taken from Table S2 of Butcher et al.2017, American Journal of Human Genetics, dx.doi.org/10.1016/j.ajhg.2017.04.004)
