## Supplementary material for "Growth disrupting mutations in epigenetic regulatory molecules are associated with abnormalities of epigenetic aging"

### Table of Contents - Supplemental Figures


- Supplemental Figure 1 – Plots of methylation values in three differentially methylated positions associated with genes
- Supplemental Figure 2 – Scatter plot examining the effect of blood composition to methylation beta values
- Supplemental Figure 3 - Example of the top ranking DMR just upstream from the gene *ANXA2R*
- Supplemental Figure 4 – Size distribution of differentially methylated regions
- Supplemental Figure 5 – Position of differentially methylated positions relative to transcripts and CpG islands

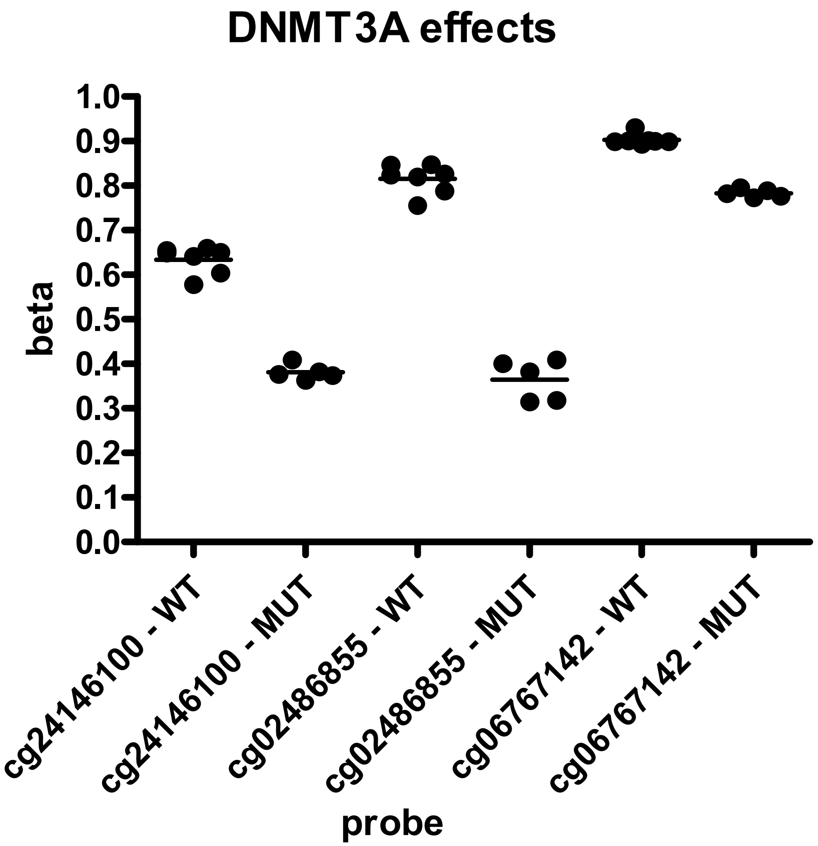


**Supplemental Figure 1** – Plots of methylation beta values shown for Amish *DNMT3A* wildtype (WT) and c.2312G>A; p.(Arg771Gln) variant carriers (MUT) for top ranking probes associated with genes: cg2416100 (associated with DOCK9), cg02486855 (associated with SMAD3) and cg06767142 (associated with MACROD1)

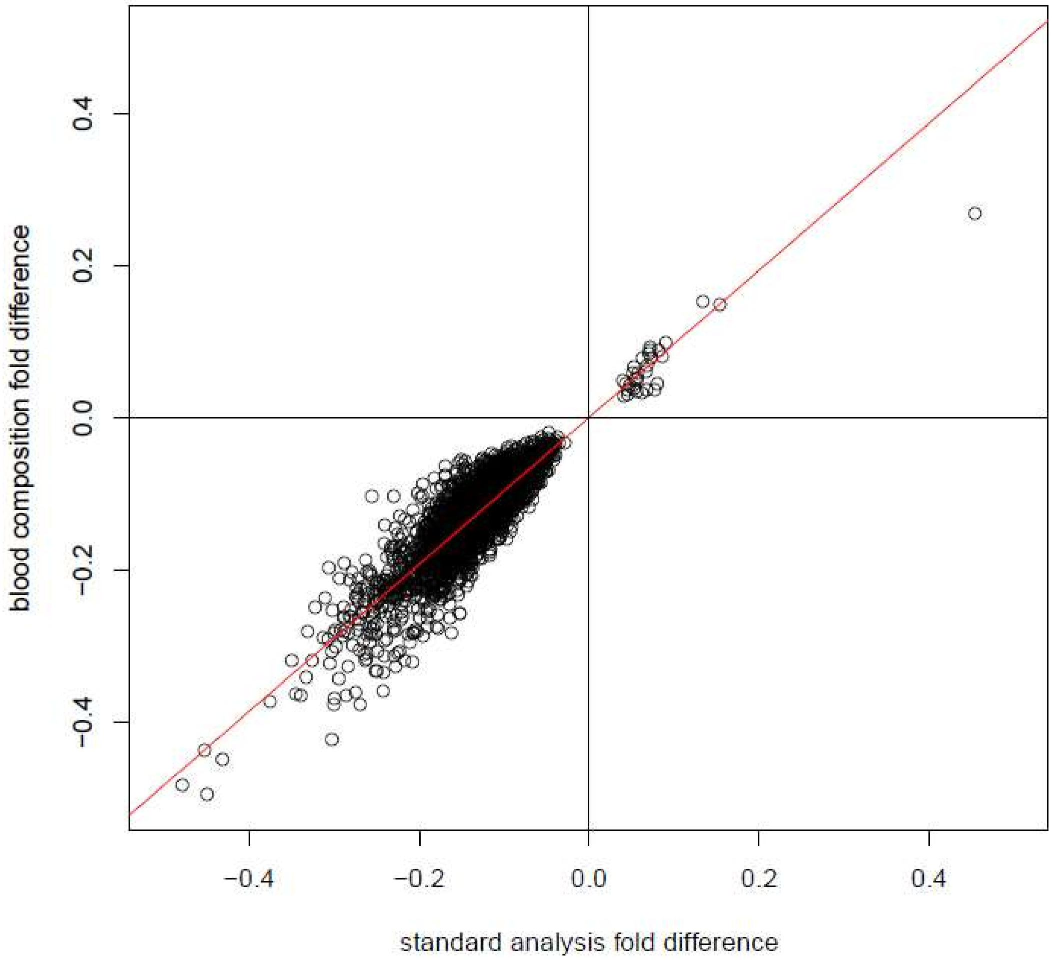


**Supplemental Figure 2** – Comparison of effect size for the 2,606 significant differentially methylated positions (DMPs) found in the standard linear model (genotype and sex) vs inclusion of blood composition (genotype, sex and predicted blood composition)

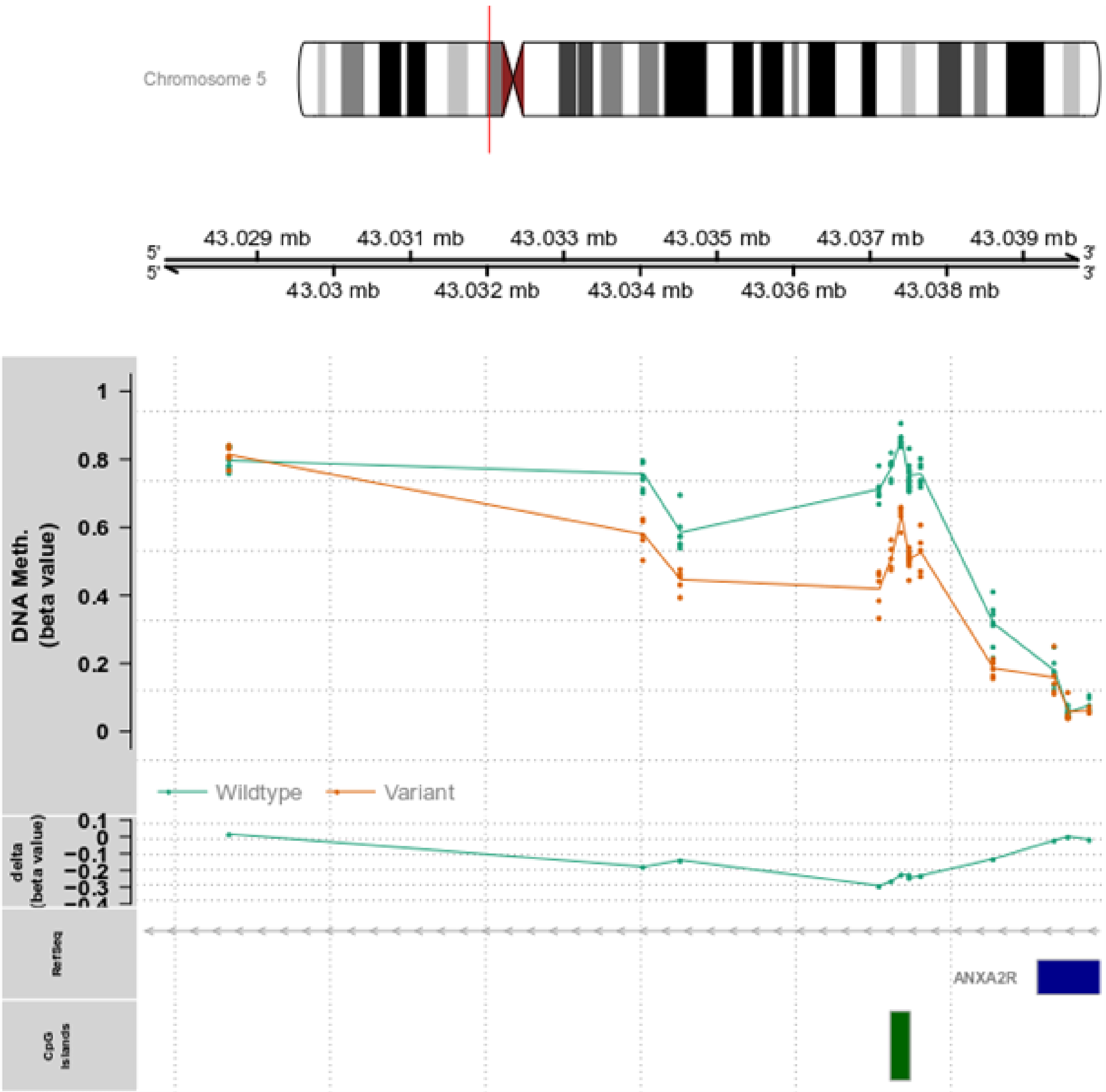


**Supplemental Figure 3** - Example of the top ranking DMR just upstream from the gene *ANXA2R*. Methylation beta values for Amish *DNMT3A* c.2312G>A; p.(Arg771Gln) variant carriers (orange) are shown together with wildtype family members (green) and their delta(beta) value difference (change in methylation) in the lower row

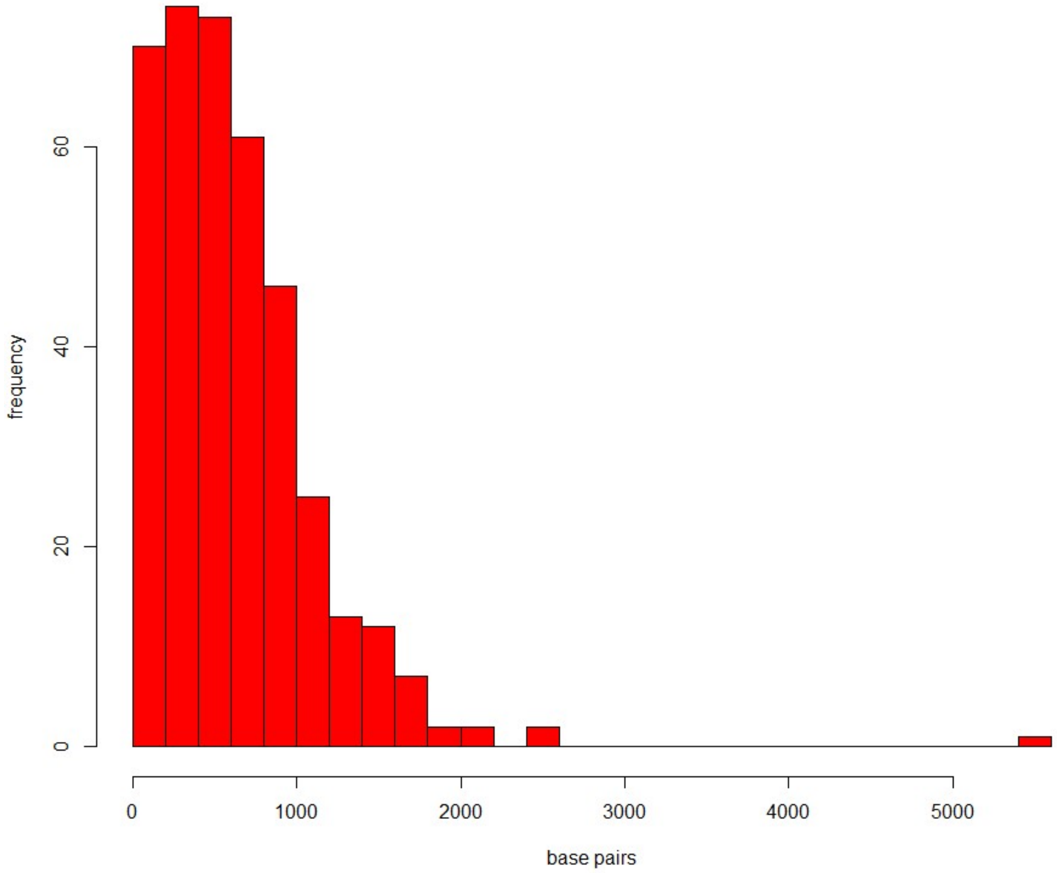


**Supplemental Figure 4** – Size distribution (in base pairs) of differentially methylated regions (DMRs) in Amish *DNMT3A* c.2312G>A; p.(Arg771Gln) variant carriers vs wildtype family members, detected on the Illumina 450k methylation beadchip array

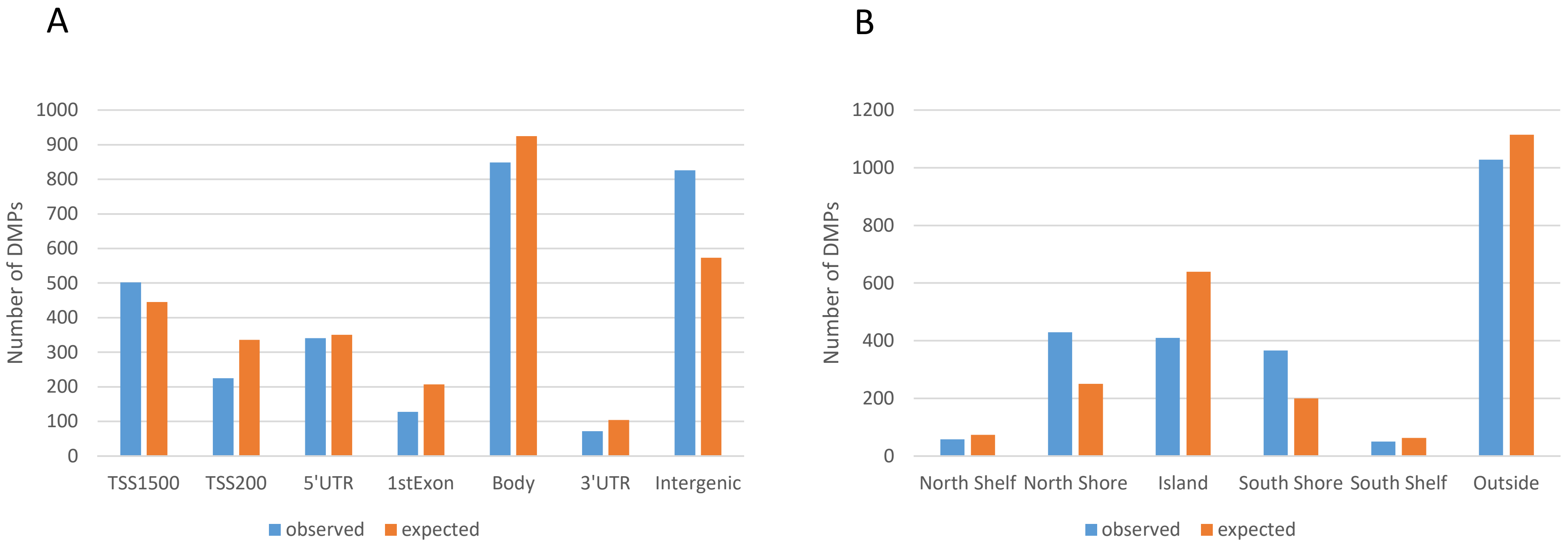


**Supplemental Figure 5** – Barchart showing the occurrence of differentially methylated positions (DMPs) annotated to regions around the transcript (A) and CpG Islands (B). The bars indicate the observed (orange) and expected number (blue) of DMPs found in Amish *DNMT3A* c.2312G>A; p.(Arg771Gln) variant carriers.

Annotation Abbreviations (from Illumina supplied 450k methylation beadchip annotation): TSS1500 = 1500bp upstream of the transcriptional start site, TSS200=200bp upstream of the transcriptional start site/the promoter, 5'UTR=5' Untranslated region, 3'UTR=3' Untranslated region, 1stExon=First exon of the transcript, Body=probes which lie within other exons or introns of the transcript. Island = CpG Island, Shore=region upto 2kb from CpG Island, Shelf=2-4kb from CpG Island, North=upstream, South=downstream, Outside=probes outside of the CpG Island defined regions.
