## Supplementary material for "Growth disrupting mutations in epigenetic regulatory molecules are associated with abnormalities of epigenetic aging"

GREAT version 3.0.0 current (02/15/2015 to now) ▾

### ⊕ Job Description

### ⊖ Region-Gene Association Graphs

*What do these graphs illustrate?*

**Number of associated genes per region**

Download as PDF.

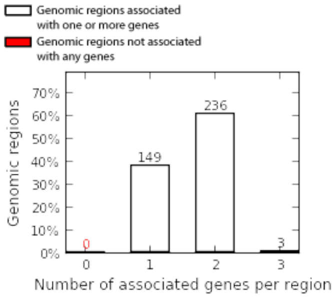


**Binned by orientation and distance to TSS**

Download as PDF.

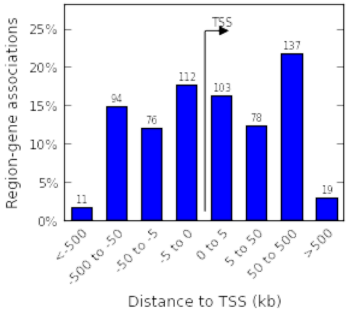


**Binned by absolute distance to TSS**

Download as PDF.

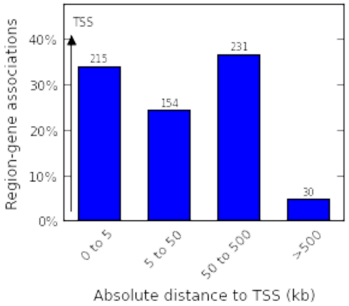


### ⊕ Global Controls

Global Export ▾ *Which data is exported by each option?*

### ⊖ GO Molecular Function (3 terms)

Global controls

*Table controls:* Export ▾ Shown top rows in this table: 20 Set Term annotation count: Min: 1 Max: Inf Set Visualize this table: [select one] ▾

| Term Name | Term ID | Binom Rank | Binom Raw P-Value | Binom Bonferroni P-Value | *Binom FDR Q-Val* | Binom Fold Enrichment | Binom Expected | Binom Observed Region Hits | Binom Genome Fraction | Binom Region Set Coverage | Hyper Rank | Hyper Raw P-Value | Hyper Bonferroni P-Value | *Hyper FDR Q-Val* | Hyper Fold Enrichment | Hyper Expected | Hyper Observed Gene Hits | Hyper Total Genes | Hyper Gene Set Coverage | Hyper Term Gene Coverage |
|---|---|---|---|---|---|---|---|---|---|---|---|---|---|---|---|---|---|---|---|---|
| **sequence-specific DNA binding** | GO:0043565 | 2 | **5.7476e-10** | **2.1197e-6** | **1.0598e-6** | 2.0959 | 36.2619 | 76 | 0.0935 | 19.59% | 2 | **4.7470e-17** | **1.7507e-13** | **8.7535e-14** | 3.1263 | 21.7511 | 68 | 697 | 12.08% | 9.76% |
| **HMG box domain binding** | GO:0071837 | 5 | **8.0649e-7** | **2.9743e-3** | **5.9486e-4** | 7.9439 | 1.2588 | 10 | 0.0032 | 2.58% | 9 | **1.7248e-5** | **6.3610e-2** | **7.0678e-3** | 10.1193 | 0.5929 | 6 | 19 | 1.07% | 31.58% |
| **regulatory region DNA binding** | GO:0000975 | 6 | **7.9757e-6** | **2.9414e-2** | **4.9024e-3** | 2.0768 | 20.2232 | 42 | 0.0521 | 10.82% | 5 | **5.4813e-8** | **2.0215e-4** | **4.0430e-5** | 2.8814 | 11.4529 | 33 | 367 | 5.86% | 8.99% |

The test set of 388 genomic regions picked 563 (3%) of all 18,041 genes.
*GO Molecular Function* has 3,688 terms covering 15,090 (84%) of all 18,041 genes, and 189,388 term - gene associations.
3,688 ontology terms (100%) were tested using an annotation count range of [1, Inf].

### ⊖ GO Biological Process (17 terms)

Global controls

*Table controls:* Export ▾ Shown top rows in this table: 20 Set Term annotation count: Min: 1 Max: Inf Set Visualize this table: [select one] ▾

| Term Name | Term ID | Binom Rank | Binom Raw P-Value | Binom Bonferroni P-Value | *Binom FDR Q-Val* | Binom Fold Enrichment | Binom Expected | Binom Observed Region Hits | Binom Genome Fraction | Binom Region Set Coverage | Hyper Rank | Hyper Raw P-Value | Hyper Bonferroni P-Value | *Hyper FDR Q-Val* | Hyper Fold Enrichment | Hyper Expected | Hyper Observed Gene Hits | Hyper Total Genes | Hyper Gene Set Coverage | Hyper Term Gene Coverage |
|---|---|---|---|---|---|---|---|---|---|---|---|---|---|---|---|---|---|---|---|---|
| **skeletal system development** | GO:0001501 | 1 | **8.8312e-11** | **9.2198e-7** | **9.2198e-7** | 2.5437 | 22.8016 | 58 | 0.0588 | 14.95% | 2 | **2.9408e-16** | **3.0702e-12** | **1.5351e-12** | 3.8962 | 12.5763 | 49 | 403 | 8.70% | 12.16% |

| Name | ID | Rank | P-Value | P-Value | *Val* | Enrichment | Expected | Region Hits | Fraction | Set Coverage | Rank | P-Value | P-Value | *Val* | Enrichment | Expected | Gene Hits | Genes | Coverage | Gene Coverage |
|---|---|---|---|---|---|---|---|---|---|---|---|---|---|---|---|---|---|---|---|---|
| **embryonic morphogenesis** | GO:0048598 | 2 | **5.2786e-10** | **5.5108e-6** | **2.7554e-6** | 2.2762 | 28.5569 | 65 | 0.0736 | 16.75% | 4 | **2.8600e-15** | **2.9858e-11** | **7.4645e-12** | 3.4063 | 15.8530 | 54 | 508 | 9.59% | 10.63% |
| **anterior/posterior pattern specification** | GO:0009952 | 3 | **6.9152e-10** | **7.2195e-6** | **2.4065e-6** | 3.3730 | 10.3765 | 35 | 0.0267 | 9.02% | 18 | **2.7401e-12** | **2.8607e-8** | **1.5893e-9** | 4.7656 | 6.0853 | 29 | 195 | 5.15% | 14.87% |
| **embryonic organ development** | GO:0048568 | 5 | **1.8247e-9** | **1.9050e-5** | **3.8101e-6** | 2.4148 | 22.7760 | 55 | 0.0587 | 14.18% | 20 | **3.1036e-12** | **3.2401e-8** | **1.6201e-9** | 3.4333 | 12.2330 | 42 | 392 | 7.46% | 10.71% |
| **embryonic organ morphogenesis** | GO:0048562 | 6 | **5.4845e-9** | **5.7258e-5** | **9.5431e-6** | 2.7034 | 15.9061 | 43 | 0.0410 | 11.08% | 25 | **1.4103e-11** | **1.4723e-7** | **5.8893e-9** | 3.9754 | 8.3010 | 33 | 266 | 5.86% | 12.41% |
| **regionalization** | GO:0003002 | 9 | **4.0631e-8** | **4.2419e-4** | **4.7132e-5** | 2.5182 | 17.0755 | 43 | 0.0440 | 11.08% | 30 | **1.9585e-11** | **2.0447e-7** | **6.8157e-9** | 3.7385 | 9.3620 | 35 | 300 | 6.22% | 11.67% |
| **pattern specification process** | GO:0007389 | 13 | **1.6930e-7** | **1.7675e-3** | **1.3596e-4** | 2.1457 | 24.7002 | 53 | 0.0637 | 13.66% | 12 | **7.0261e-13** | **7.3353e-9** | **6.1127e-10** | 3.4009 | 13.2316 | 45 | 424 | 7.99% | 10.61% |
| **skeletal system morphogenesis** | GO:0048705 | 28 | **9.7855e-7** | **1.0216e-2** | **3.6486e-4** | 2.7370 | 10.9607 | 30 | 0.0282 | 7.73% | 23 | **9.3387e-12** | **9.7496e-8** | **4.2390e-9** | 4.6976 | 5.9605 | 28 | 191 | 4.97% | 14.66% |
| **embryonic skeletal system morphogenesis** | GO:0048704 | 33 | **2.2968e-6** | **2.3979e-2** | **7.2664e-4** | 3.6094 | 5.2640 | 19 | 0.0136 | 4.90% | 47 | **1.9138e-10** | **1.9980e-6** | **4.2511e-8** | 6.5545 | 2.7462 | 18 | 88 | 3.20% | 20.45% |
| **embryonic skeletal system development** | GO:0048706 | 41 | **4.5261e-6** | **4.7252e-2** | **1.1525e-3** | 2.9918 | 7.6876 | 23 | 0.0198 | 5.93% | 15 | **1.3688e-12** | **1.4290e-8** | **9.5269e-10** | 6.2993 | 3.6512 | 23 | 117 | 4.09% | 19.66% |
| **ossification** | GO:0001503 | 52 | **1.1910e-5** | **1.2434e-1** | **2.3912e-3** | 2.6096 | 9.9634 | 26 | 0.0257 | 6.70% | 92 | **3.7858e-6** | **3.9524e-2** | **4.2961e-4** | 3.2532 | 6.1477 | 20 | 197 | 3.55% | 10.15% |
| **muscle organ development** | GO:0007517 | 67 | **5.3845e-5** | **5.6215e-1** | **8.3903e-3** | 2.2155 | 13.5410 | 30 | 0.0349 | 7.73% | 111 | **2.9925e-5** | **3.1241e-1** | **2.8145e-3** | 2.6704 | 8.2386 | 22 | 264 | 3.91% | 8.33% |
| **chondroblast differentiation** | GO:0060591 | 69 | **6.1310e-5** | **6.4007e-1** | **9.2764e-3** | 41.0115 | 0.0732 | 3 | 0.0002 | 0.77% | 131 | **1.1812e-4** | **1.0000** | **9.4136e-3** | 24.0333 | 0.1248 | 3 | 4 | 0.53% | 75.00% |
| **negative regulation of myeloid cell differentiation** | GO:0045638 | 83 | **1.5678e-4** | **1.0000** | **1.9720e-2** | 3.6344 | 3.3018 | 12 | 0.0085 | 3.09% | 143 | **1.5114e-4** | **1.0000** | **1.1034e-2** | 4.1083 | 2.4341 | 10 | 78 | 1.78% | 12.82% |
| **positive regulation of steroid biosynthetic process** | GO:0010893 | 89 | **2.3134e-4** | **1.0000** | **2.7137e-2** | 9.3321 | 0.5358 | 5 | 0.0014 | 1.29% | 158 | **3.8036e-4** | **1.0000** | **2.5133e-2** | 10.6815 | 0.3745 | 4 | 12 | 0.71% | 33.33% |
| **osteoblast development** | GO:0002076 | 94 | **3.2381e-4** | **1.0000** | **3.5963e-2** | 5.5549 | 1.2601 | 7 | 0.0032 | 1.80% | 148 | **2.3507e-4** | **1.0000** | **1.6582e-2** | 8.4327 | 0.5929 | 5 | 19 | 0.89% | 26.32% |
| **artery morphogenesis** | GO:0048844 | 103 | **4.8435e-4** | **1.0000** | **4.9093e-2** | 3.4182 | 3.2181 | 11 | 0.0083 | 2.84% | 172 | **6.7334e-4** | **1.0000** | **4.0870e-2** | 4.6731 | 1.4979 | 7 | 48 | 1.24% | 14.58% |

The test set of 388 genomic regions picked 563 (3%) of all 18,041 genes.
*GO Biological Process* has 10,440 terms covering 15,441 (86%) of all 18,041 genes, and 950,065 term - gene associations.
10,440 ontology terms (100%) were tested using an annotation count range of [1, Inf].

### ⊖ GO Cellular Component (1 term)

Global controls

*Table controls:* Export ▾ Shown top rows in this table: 20 Set Term annotation count: Min: 1 Max: Inf Set Visualize this table: NEW [select one] ▾

| Term Name | Term ID | Binom Rank | Binom Raw P-Value | Binom Bonferroni P-Value | *Binom FDR Q-Val* | Binom Fold Enrichment | Binom Expected | Binom Observed Region Hits | Binom Genome Fraction | Binom Region Set Coverage | Hyper Rank | Hyper Raw P-Value | Hyper Bonferroni P-Value | *Hyper FDR Q-Val* | Hyper Fold Enrichment | Hyper Expected | Hyper Observed Gene Hits | Hyper Total Genes | Hyper Gene Set Coverage | Hyper Term Gene Coverage |
|---|---|---|---|---|---|---|---|---|---|---|---|---|---|---|---|---|---|---|---|---|
| **PcG protein complex** | GO:0031519 | 4 | **2.6739e-5** | **3.3825e-2** | **8.4563e-3** | 5.9780 | 1.5055 | 9 | 0.0039 | 2.32% | 2 | **2.2406e-5** | **2.8344e-2** | **1.4172e-2** | 6.5732 | 1.2171 | 8 | 39 | 1.42% | 20.51% |

The test set of 388 genomic regions picked 563 (3%) of all 18,041 genes.
*GO Cellular Component* has 1,265 terms covering 16,807 (93%) of all 18,041 genes, and 277,316 term - gene associations.
1,265 ontology terms (100%) were tested using an annotation count range of [1, Inf].

*Table controls:* Export ▾ Shown top rows in this table: 20 Set Term annotation count: Min: 1 Max: Inf Set Visualize this table: [select one] ▾

| Term Name | Term ID | Binom Rank | Binom Raw P-Value | Binom Bonferroni P-Value | *Binom FDR Q-Val* | Binom Fold Enrichment | Binom Expected | Binom Observed Region Hits | Binom Genome Fraction | Binom Region Set Coverage | Hyper Rank | Hyper Raw P-Value | Hyper Bonferroni P-Value | *Hyper FDR Q-Val* | Hyper Fold Enrichment | Hyper Expected | Hyper Observed Gene Hits | Hyper Total Genes | Hyper Gene Set Coverage | Hyper Term Gene Coverage |
|---|---|---|---|---|---|---|---|---|---|---|---|---|---|---|---|---|---|---|---|---|
| thoracic vertebral transformation | MP:0004618 | 1 | **4.4458e-14** | **3.5202e-10** | **3.5202e-10** | 11.9222 | 1.5098 | 18 | 0.0039 | 4.64% | 17 | **6.0629e-11** | **4.8006e-7** | **2.8239e-8** | 8.9012 | 1.6852 | 15 | 54 | 2.66% | 27.78% |
| abnormal sternebra morphology | MP:0004322 | 2 | **2.7539e-11** | **2.1806e-7** | **1.0903e-7** | 6.5333 | 3.2143 | 21 | 0.0083 | 5.41% | 22 | **2.4197e-10** | **1.9159e-6** | **8.7088e-8** | 8.1469 | 1.8412 | 15 | 59 | 2.66% | 25.42% |
| abnormal rib development | MP:0002823 | 3 | **2.8995e-11** | **2.2958e-7** | **7.6527e-8** | 11.9491 | 1.1716 | 14 | 0.0030 | 3.61% | 126 | **4.7384e-5** | **3.7518e-1** | **2.9776e-3** | 7.0097 | 0.9986 | 7 | 32 | 1.24% | 21.88% |
| abnormal cervical vertebrae morphology | MP:0003048 | 4 | **4.5629e-10** | **3.6129e-6** | **9.0323e-7** | 4.4541 | 5.8374 | 26 | 0.0150 | 6.70% | 12 | **1.0731e-11** | **8.4969e-8** | **7.0808e-9** | 6.0254 | 3.6512 | 22 | 117 | 3.91% | 18.80% |
| abnormal vertebrae morphology | MP:0000137 | 5 | **5.8125e-10** | **4.6024e-6** | **9.2047e-7** | 2.6774 | 18.3013 | 49 | 0.0472 | 12.63% | 9 | **3.5959e-12** | **2.8472e-8** | **3.1636e-9** | 3.5506 | 11.2656 | 40 | 361 | 7.10% | 11.08% |
| vertebral transformation | MP:0003036 | 6 | **7.7012e-10** | **6.0978e-6** | **1.0163e-6** | 5.4186 | 3.8756 | 21 | 0.0100 | 5.41% | 35 | **3.8747e-9** | **3.0680e-5** | **8.7657e-7** | 5.4933 | 3.2767 | 18 | 105 | 3.20% | 17.14% |
| abnormal rib morphology | MP:0000150 | 7 | **8.4592e-10** | **6.6980e-6** | **9.5686e-7** | 3.0860 | 12.6378 | 39 | 0.0326 | 10.05% | 30 | **1.1165e-9** | **8.8401e-6** | **2.9467e-7** | 3.7321 | 7.7705 | 29 | 249 | 5.15% | 11.65% |
| abnormal thoracic cage morphology | MP:0004624 | 8 | **9.3369e-10** | **7.3929e-6** | **9.2412e-7** | 2.6059 | 19.1869 | 50 | 0.0495 | 12.89% | 8 | **2.5109e-12** | **1.9881e-8** | **2.4851e-9** | 3.6649 | 10.6415 | 39 | 341 | 6.93% | 11.44% |
| abnormal sternum morphology | MP:0000157 | 9 | **1.1460e-9** | **9.0737e-6** | **1.0082e-6** | 3.0522 | 12.7775 | 39 | 0.0329 | 10.05% | 13 | **1.1043e-11** | **8.7435e-8** | **6.7258e-9** | 4.5111 | 6.4286 | 29 | 206 | 5.15% | 14.08% |
| abnormal presacral vertebrae morphology | MP:0000459 | 10 | **1.4313e-9** | **1.1333e-5** | **1.1333e-6** | 3.2783 | 10.6764 | 35 | 0.0275 | 9.02% | 14 | **1.2473e-11** | **9.8759e-8** | **7.0542e-9** | 4.4893 | 6.4598 | 29 | 207 | 5.15% | 14.01% |
| abnormal rib attachment | MP:0004625 | 11 | **3.4050e-9** | **2.6961e-5** | **2.4510e-6** | 4.7393 | 4.6421 | 22 | 0.0120 | 5.67% | 28 | **7.1791e-10** | **5.6844e-6** | **2.0302e-7** | 6.0716 | 2.9646 | 18 | 95 | 3.20% | 18.95% |
| abnormal pectoral girdle bone morphology | MP:0004508 | 12 | **3.7139e-9** | **2.9407e-5** | **2.4506e-6** | 2.7822 | 15.0958 | 42 | 0.0389 | 10.82% | 7 | **1.8438e-12** | **1.4599e-8** | **2.0855e-9** | 4.4009 | 7.2712 | 32 | 233 | 5.68% | 13.73% |
| cervical vertebral transformation | MP:0004615 | 14 | **6.3805e-9** | **5.0521e-5** | **3.6086e-6** | 8.6979 | 1.4946 | 13 | 0.0039 | 3.35% | 40 | **8.5275e-9** | **6.7521e-5** | **1.6880e-6** | 8.5452 | 1.4043 | 12 | 45 | 2.13% | 26.67% |
| abnormal appendicular skeleton morphology | MP:0009250 | 15 | **7.4373e-9** | **5.8888e-5** | **3.9259e-6** | 2.0960 | 31.9653 | 67 | 0.0824 | 17.27% | 3 | **6.6323e-14** | **5.2514e-10** | **1.7505e-10** | 3.0780 | 18.1935 | 56 | 583 | 9.95% | 9.61% |
| increased rib number | MP:0000480 | 16 | **7.8694e-9** | **6.2310e-5** | **3.8944e-6** | 6.9950 | 2.1444 | 15 | 0.0055 | 3.87% | 47 | **9.6083e-8** | **7.6079e-4** | **1.6187e-5** | 7.8331 | 1.4043 | 11 | 45 | 1.95% | 24.44% |
| abnormal kidney venous blood vessel morphology | MP:0011311 | 17 | **9.0416e-9** | **7.1591e-5** | **4.2113e-6** | 42.7745 | 0.1403 | 6 | 0.0004 | 1.55% | 156 | **1.1812e-4** | **9.3528e-1** | **5.9954e-3** | 24.0333 | 0.1248 | 3 | 4 | 0.53% | 75.00% |
| small xiphoid process | MP:0004680 | 18 | **1.2869e-8** | **1.0189e-4** | **5.6607e-6** | 40.2817 | 0.1490 | 6 | 0.0004 | 1.55% | 134 | **5.9436e-5** | **4.7061e-1** | **3.5120e-3** | 16.0222 | 0.2497 | 4 | 8 | 0.71% | 50.00% |
| abnormal vertebral column morphology | MP:0004703 | 19 | **5.3617e-8** | **4.2454e-4** | **2.2344e-5** | 2.1109 | 27.9499 | 59 | 0.0720 | 15.21% | 20 | **8.6317e-11** | **6.8346e-7** | **3.4173e-8** | 2.7939 | 17.5382 | 49 | 562 | 8.70% | 8.72% |
| abnormal cervical atlas morphology | MP:0004607 | 20 | **5.6137e-8** | **4.4450e-4** | **2.2225e-5** | 6.5404 | 2.1405 | 14 | 0.0055 | 3.61% | 71 | **1.8171e-6** | **1.4388e-2** | **2.0264e-4** | 6.6759 | 1.4979 | 10 | 48 | 1.78% | 20.83% |
| abnormal vertebral arch morphology | MP:0004599 | 21 | **6.7945e-8** | **5.3799e-4** | **2.5618e-5** | 4.1628 | 5.0446 | 21 | 0.0130 | 5.41% | 72 | **2.0730e-6** | **1.6414e-2** | **2.2797e-4** | 4.5778 | 3.0583 | 14 | 98 | 2.49% | 14.29% |

The test set of 388 genomic regions picked 563 (3%) of all 18,041 genes.
*Mouse Phenotype* has 7,918 terms covering 7,524 (42%) of all 18,041 genes, and 541,054 term - gene associations.
7,918 ontology terms (100%) were tested using an annotation count range of [1, Inf].

### ⊖ Human Phenotype (no terms)

Global controls

| Term Name | Term ID | Binom Rank | Binom Raw P-Value | Binom Bonferroni P-Value | *Binom FDR Q-Val* | Binom Fold Enrichment | Binom Expected | Binom Observed Region Hits | Binom Genome Fraction | Binom Region Set Coverage | Hyper Rank | Hyper Raw P-Value | Hyper Bonferroni P-Value | *Hyper FDR Q-Val* | Hyper Fold Enrichment | Hyper Expected | Hyper Observed Gene Hits | Hyper Total Genes | Hyper Gene Set Coverage | Hyper Term Gene Coverage |
|---|---|---|---|---|---|---|---|---|---|---|---|---|---|---|---|---|---|---|---|---|

No results meet your chosen criteria.

The test set of 388 genomic regions picked 563 (3%) of all 18,041 genes.
*Human Phenotype* has 6,146 terms covering 2,822 (16%) of all 18,041 genes, and 244,804 term - gene associations.
6,146 ontology terms (100%) were tested using an annotation count range of [1, Inf].

### Disease Ontology (no terms)

Global controls

*Table controls:* Export ▾ Shown top rows in this table: 20 Set Term annotation count: Min: 1 Max: Inf Set Visualize this table: NEW [select one] ▾

| Term Name | Term ID | Binom Rank | Binom Raw P-Value | Binom Bonferroni P-Value | *Binom FDR Q-Val* | Binom Fold Enrichment | Binom Expected | Binom Observed Region Hits | Binom Genome Fraction | Binom Region Set Coverage | Hyper Rank | Hyper Raw P-Value | Hyper Bonferroni P-Value | *Hyper FDR Q-Val* | Hyper Fold Enrichment | Hyper Expected | Hyper Observed Gene Hits | Hyper Total Genes | Hyper Gene Set Coverage | Hyper Term Gene Coverage |
|---|---|---|---|---|---|---|---|---|---|---|---|---|---|---|---|---|---|---|---|---|

No results meet your chosen criteria.

The test set of 388 genomic regions picked 563 (3%) of all 18,041 genes.
*Disease Ontology* has 2,235 terms covering 7,886 (44%) of all 18,041 genes, and 232,324 term - gene associations.
2,235 ontology terms (100%) were tested using an annotation count range of [1, Inf].

### MSigDB Cancer Neighborhood (no terms)

Global controls

*Table controls:* Export ▾ Shown top rows in this table: 20 Set Term annotation count: Min: 1 Max: Inf Set Visualize this table: NEW [select one] ▾

| Term Name | Term ID | Binom Rank | Binom Raw P-Value | Binom Bonferroni P-Value | *Binom FDR Q-Val* | Binom Fold Enrichment | Binom Expected | Binom Observed Region Hits | Binom Genome Fraction | Binom Region Set Coverage | Hyper Rank | Hyper Raw P-Value | Hyper Bonferroni P-Value | *Hyper FDR Q-Val* | Hyper Fold Enrichment | Hyper Expected | Hyper Observed Gene Hits | Hyper Total Genes | Hyper Gene Set Coverage | Hyper Term Gene Coverage |
|---|---|---|---|---|---|---|---|---|---|---|---|---|---|---|---|---|---|---|---|---|

No results meet your chosen criteria.

The test set of 388 genomic regions picked 563 (3%) of all 18,041 genes.
*MSigDB Cancer Neighborhood* has 427 terms covering 4,758 (26%) of all 18,041 genes, and 42,051 term - gene associations.
427 ontology terms (100%) were tested using an annotation count range of [1, Inf].

### Placenta Disorders (no terms)

Global controls

*Table controls:* Export ▾ Shown top rows in this table: 20 Set Term annotation count: Min: 1 Max: Inf Set Visualize this table: NEW [select one] ▾

| Term Name | Term ID | Binom Rank | Binom Raw P-Value | Binom Bonferroni P-Value | *Binom FDR Q-Val* | Binom Fold Enrichment | Binom Expected | Binom Observed Region Hits | Binom Genome Fraction | Binom Region Set Coverage | Hyper Rank | Hyper Raw P-Value | Hyper Bonferroni P-Value | *Hyper FDR Q-Val* | Hyper Fold Enrichment | Hyper Expected | Hyper Observed Gene Hits | Hyper Total Genes | Hyper Gene Set Coverage | Hyper Term Gene Coverage |
|---|---|---|---|---|---|---|---|---|---|---|---|---|---|---|---|---|---|---|---|---|

No results meet your chosen criteria.

The test set of 388 genomic regions picked 563 (3%) of all 18,041 genes.
*Placenta Disorders* has 3 terms covering 691 (4%) of all 18,041 genes, and 831 term - gene associations.
3 ontology terms (100%) were tested using an annotation count range of [1, Inf].

### PANTHER Pathway (no terms)

Global controls

| Term Name | Term ID | Binom Rank | Binom Raw P-Value | Binom Bonferroni P-Value | *Binom FDR Q-Val* | Binom Fold Enrichment | Binom Expected | Binom Observed Region Hits | Binom Genome Fraction | Binom Region Set Coverage | Hyper Rank | Hyper Raw P-Value | Hyper Bonferroni P-Value | *Hyper FDR Q-Val* | Hyper Fold Enrichment | Hyper Expected | Hyper Observed Gene Hits | Hyper Total Genes | Hyper Gene Set Coverage | Hyper Term Gene Coverage |
|---|---|---|---|---|---|---|---|---|---|---|---|---|---|---|---|---|---|---|---|---|

No results meet your chosen criteria.

The test set of 388 genomic regions picked 563 (3%) of all 18,041 genes.
*PANTHER Pathway* has 152 terms covering 2,197 (12%) of all 18,041 genes, and 5,065 term - gene associations.
152 ontology terms (100%) were tested using an annotation count range of [1, Inf].

### BioCyc Pathway (no terms)

Global controls

*Table controls:* Export ▾ Shown top rows in this table: 20 Set Term annotation count: Min: 1 Max: Inf Set Visualize this table: [select one] ▾

| Term Name | Term ID | Binom Rank | Binom Raw P-Value | Binom Bonferroni P-Value | *Binom FDR Q-Val* | Binom Fold Enrichment | Binom Expected | Binom Observed Region Hits | Binom Genome Fraction | Binom Region Set Coverage | Hyper Rank | Hyper Raw P-Value | Hyper Bonferroni P-Value | *Hyper FDR Q-Val* | Hyper Fold Enrichment | Hyper Expected | Hyper Observed Gene Hits | Hyper Total Genes | Hyper Gene Set Coverage | Hyper Term Gene Coverage |
|---|---|---|---|---|---|---|---|---|---|---|---|---|---|---|---|---|---|---|---|---|

No results meet your chosen criteria.

The test set of 388 genomic regions picked 563 (3%) of all 18,041 genes.
*BioCyc Pathway* has 332 terms covering 1,034 (6%) of all 18,041 genes, and 2,524 term - gene associations.
332 ontology terms (100%) were tested using an annotation count range of [1, Inf].

### MSigDB Pathway (no terms)

Global controls

*Table controls:* Export ▾ Shown top rows in this table: 20 Set Term annotation count: Min: 1 Max: Inf Set Visualize this table: [select one] ▾

| Term Name | Term ID | Binom Rank | Binom Raw P-Value | Binom Bonferroni P-Value | *Binom FDR Q-Val* | Binom Fold Enrichment | Binom Expected | Binom Observed Region Hits | Binom Genome Fraction | Binom Region Set Coverage | Hyper Rank | Hyper Raw P-Value | Hyper Bonferroni P-Value | *Hyper FDR Q-Val* | Hyper Fold Enrichment | Hyper Expected | Hyper Observed Gene Hits | Hyper Total Genes | Hyper Gene Set Coverage | Hyper Term Gene Coverage |
|---|---|---|---|---|---|---|---|---|---|---|---|---|---|---|---|---|---|---|---|---|

No results meet your chosen criteria.

The test set of 388 genomic regions picked 563 (3%) of all 18,041 genes.
*MSigDB Pathway* has 1,320 terms covering 8,117 (45%) of all 18,041 genes, and 62,449 term - gene associations.
1,320 ontology terms (100%) were tested using an annotation count range of [1, Inf].

Term Name Term ID Binom Rank Binom Raw P-Value Binom Bonferroni P-Value Binom FDR Q-Val Binom Fold Enrichment Binom Expected Binom Observed Region Hits Binom Genome Fraction Binom Region Set Coverage Hyper Rank Hyper Raw P-Value Hyper Bonferroni P-Value Hyper FDR Q-Val Hyper Fold Enrichment Hyper Expected Hyper Observed Gene Hits Hyper Total Genes Hyper Gene Set Coverage Hyper Term Gene Coverage

### MGI Expression: Detected (20+ terms)

Global controls

*Table controls:* Export ▾ Shown top rows in this table: 20 Set Term annotation count: Min: 1 Max: Inf Set Visualize this table: [select one] ▾

| Term Name | Term ID | Binom Rank | Binom Raw P-Value | Binom Bonferroni P-Value | *Binom FDR Q-Val* | Binom Fold Enrichment | Binom Expected | Binom Observed Region Hits | Binom Genome Fraction | Binom Region Set Coverage | Hyper Rank | Hyper Raw P-Value | Hyper Bonferroni P-Value | *Hyper FDR Q-Val* | Hyper Fold Enrichment | Hyper Expected | Hyper Observed Gene Hits | Hyper Total Genes | Hyper Gene Set Coverage | Hyper Term Gene Coverage |
|---|---|---|---|---|---|---|---|---|---|---|---|---|---|---|---|---|---|---|---|---|
| **TS19_T1 vertebral cartilage condensation** | 13390 | 1 | **6.3449e-17** | **5.9243e-13** | **5.9243e-13** | 422.5308 | 0.0166 | 7 | 4.2698e-5 | 1.80% | 127 | **2.3884e-8** | **2.2300e-4** | **1.7559e-6** | 24.0333 | 0.2497 | 6 | 8 | 1.07% | 75.00% |
| **TS19_T2 vertebral cartilage condensation** | 13395 | 2 | **1.2054e-16** | **1.1255e-12** | **5.6273e-13** | 385.4422 | 0.0182 | 7 | 4.6807e-5 | 1.80% | 148 | **6.9756e-8** | **6.5131e-4** | **4.4008e-6** | 21.3629 | 0.2809 | 6 | 9 | 1.07% | 66.67% |
| **TS19_T3 vertebral cartilage condensation** | 13400 | 2 | **1.2054e-16** | **1.1255e-12** | **5.6273e-13** | 385.4422 | 0.0182 | 7 | 4.6807e-5 | 1.80% | 148 | **6.9756e-8** | **6.5131e-4** | **4.4008e-6** | 21.3629 | 0.2809 | 6 | 9 | 1.07% | 66.67% |

| Name | ID | Rank | P-Value | P-Value | *Val* | Enrichment | Expected | Region Hits | Fraction | Set Coverage | Rank | P-Value | P-Value | *Val* | Enrichment | Expected | Gene Hits | Genes | Coverage | Gene Coverage |
|---|---|---|---|---|---|---|---|---|---|---|---|---|---|---|---|---|---|---|---|---|
| **TS19_T11 vertebral cartilage condensation** | 13450 | 4 | **1.8918e-16** | **1.7664e-12** | **4.4160e-13** | 361.3513 | 0.0194 | 7 | 4.9927e-5 | 1.80% | 92 | **3.0729e-9** | **2.8691e-5** | **3.1186e-7** | 22.4311 | 0.3121 | 7 | 10 | 1.24% | 70.00% |
| **TS19_T5 vertebral cartilage condensation** | 13410 | 4 | **1.8918e-16** | **1.7664e-12** | **4.4160e-13** | 361.3513 | 0.0194 | 7 | 4.9927e-5 | 1.80% | 92 | **3.0729e-9** | **2.8691e-5** | **3.1186e-7** | 22.4311 | 0.3121 | 7 | 10 | 1.24% | 70.00% |
| **TS19_T4 vertebral cartilage condensation** | 13405 | 4 | **1.8918e-16** | **1.7664e-12** | **4.4160e-13** | 361.3513 | 0.0194 | 7 | 4.9927e-5 | 1.80% | 92 | **3.0729e-9** | **2.8691e-5** | **3.1186e-7** | 22.4311 | 0.3121 | 7 | 10 | 1.24% | 70.00% |
| **TS19_T6 vertebral cartilage condensation** | 13430 | 4 | **1.8918e-16** | **1.7664e-12** | **4.4160e-13** | 361.3513 | 0.0194 | 7 | 4.9927e-5 | 1.80% | 92 | **3.0729e-9** | **2.8691e-5** | **3.1186e-7** | 22.4311 | 0.3121 | 7 | 10 | 1.24% | 70.00% |
| **TS19_T7 vertebral cartilage condensation** | 13434 | 4 | **1.8918e-16** | **1.7664e-12** | **4.4160e-13** | 361.3513 | 0.0194 | 7 | 4.9927e-5 | 1.80% | 92 | **3.0729e-9** | **2.8691e-5** | **3.1186e-7** | 22.4311 | 0.3121 | 7 | 10 | 1.24% | 70.00% |
| **TS19_T8 vertebral cartilage condensation** | 13438 | 4 | **1.8918e-16** | **1.7664e-12** | **4.4160e-13** | 361.3513 | 0.0194 | 7 | 4.9927e-5 | 1.80% | 92 | **3.0729e-9** | **2.8691e-5** | **3.1186e-7** | 22.4311 | 0.3121 | 7 | 10 | 1.24% | 70.00% |
| **TS19_T9 vertebral cartilage condensation** | 13442 | 4 | **1.8918e-16** | **1.7664e-12** | **4.4160e-13** | 361.3513 | 0.0194 | 7 | 4.9927e-5 | 1.80% | 92 | **3.0729e-9** | **2.8691e-5** | **3.1186e-7** | 22.4311 | 0.3121 | 7 | 10 | 1.24% | 70.00% |
| **TS19_T10 vertebral cartilage condensation** | 13446 | 4 | **1.8918e-16** | **1.7664e-12** | **4.4160e-13** | 361.3513 | 0.0194 | 7 | 4.9927e-5 | 1.80% | 92 | **3.0729e-9** | **2.8691e-5** | **3.1186e-7** | 22.4311 | 0.3121 | 7 | 10 | 1.24% | 70.00% |
| **TS19_C6 vertebral cartilage condensation** | 13368 | 12 | **2.7574e-15** | **2.5746e-11** | **2.1455e-12** | 530.8977 | 0.0113 | 6 | 2.9128e-5 | 1.55% | 174 | **1.7004e-7** | **1.5877e-3** | **9.1245e-6** | 26.7037 | 0.1872 | 5 | 6 | 0.89% | 83.33% |
| **TS19_C7 vertebral cartilage condensation** | 13373 | 13 | **1.0607e-14** | **9.9041e-11** | **7.6185e-12** | 423.9541 | 0.0142 | 6 | 3.6475e-5 | 1.55% | 211 | **5.7981e-7** | **5.4137e-3** | **2.5657e-5** | 22.8889 | 0.2184 | 5 | 7 | 0.89% | 71.43% |
| **TS19_cervical vertebral cartilage condensation** | 3984 | 14 | **1.4604e-12** | **1.3636e-8** | **9.7398e-10** | 186.0960 | 0.0322 | 6 | 8.3096e-5 | 1.55% | 282 | **6.4344e-6** | **6.0078e-2** | **2.1304e-4** | 16.0222 | 0.3121 | 5 | 10 | 0.89% | 50.00% |
| **TS26_integumental system** | 7528 | 15 | **3.5505e-10** | **3.3151e-6** | **2.2101e-7** | 3.5434 | 9.5952 | 34 | 0.0247 | 8.76% | 141 | **5.6883e-8** | **5.3112e-4** | **3.7668e-6** | 3.7412 | 6.1477 | 23 | 197 | 4.09% | 11.68% |
| **TS26_skin** | 14166 | 17 | **9.7630e-10** | **9.1157e-6** | **5.3622e-7** | 4.2928 | 6.0566 | 26 | 0.0156 | 6.70% | 180 | **2.1483e-7** | **2.0059e-3** | **1.1144e-5** | 4.2726 | 4.2129 | 18 | 135 | 3.20% | 13.33% |
| **TS13_trunk mesenchyme** | 498 | 19 | **3.0122e-9** | **2.8125e-5** | **1.4802e-6** | 3.2528 | 10.4526 | 34 | 0.0269 | 8.76% | 36 | **1.8582e-12** | **1.7350e-8** | **4.8195e-10** | 5.0125 | 5.5860 | 28 | 179 | 4.97% | 15.64% |
| **TS13_trunk paraxial mesenchyme** | 503 | 20 | **3.2320e-9** | **3.0177e-5** | **1.5089e-6** | 4.1961 | 5.9579 | 25 | 0.0154 | 6.44% | 80 | **1.4466e-9** | **1.3507e-5** | **1.6884e-7** | 5.8263 | 3.0895 | 18 | 99 | 3.20% | 18.18% |
| **TS15_limb** | 1451 | 21 | **1.1464e-8** | **1.0704e-4** | **5.0969e-6** | 2.1879 | 27.4238 | 60 | 0.0707 | 15.46% | 43 | **9.3064e-12** | **8.6893e-8** | **2.0208e-9** | 3.0612 | 15.3537 | 47 | 492 | 8.35% | 9.55% |
| **TS25_metanephros** | 8015 | 23 | **3.7148e-8** | **3.4685e-4** | **1.5081e-5** | 3.1266 | 9.9151 | 31 | 0.0256 | 7.99% | 68 | **4.6405e-10** | **4.3328e-6** | **6.3718e-8** | 4.1200 | 6.5534 | 27 | 210 | 4.80% | 12.86% |

The test set of 388 genomic regions picked 563 (3%) of all 18,041 genes.
*MGI Expression: Detected* has 9,337 terms covering 11,602 (64%) of all 18,041 genes, and 970,237 term - gene associations.
9,337 ontology terms (100%) were tested using an annotation count range of [1, Inf].

### MSigDB Perturbation (20+ terms)

Global controls

*Table controls:* Export ▾ Shown top rows in this table: 20 Set Term annotation count: Min: 1 Max: Inf Set Visualize this table: NEW [select one] ▾

| Term Name | Term ID | Binom Rank | Binom Raw P-Value | Binom Bonferroni P-Value | *Binom FDR Q-Val* | Binom Fold Enrichment | Binom Expected | Binom Observed Region Hits | Binom Genome Fraction | Binom Region Set Coverage | Hyper Rank | Hyper Raw P-Value | Hyper Bonferroni P-Value | *Hyper FDR Q-Val* | Hyper Fold Enrichment | [illegible] |
|---|---|---|---|---|---|---|---|---|---|---|---|---|---|---|---|---|
| **Genes with promoters occupied by PML-RARA fusion [GeneID=5371,5914] protein in acute promyelocytic leukemia(APL) cells NB4 and two APL primary blasts, based on Chip-seq data.** | MARTENS_BOUND_BY_PML_RARA_FUSION | 1 | **1.8373e-10** | **6.1808e-7** | **6.1808e-7** | 3.1392 | 13.0608 | 41 | 0.0337 | 10.57% | 3 | **1.6222e-9** | **5.4571e-6** | **1.8190e-6** | 2.9544 | 1 |

| Name | ID | Rank | P-Value | Bonferroni P-Value | FDR Q-Val | Fold Enrichment | Expected | Region Hits | Genome Fraction | Set Coverage | Rank | Raw P-Value | Bonferroni P-Value | FDR Q-Val | Fold Enrichment | Ex |
|---|---|---|---|---|---|---|---|---|---|---|---|---|---|---|---|---|
| **Genes with high-CpG-density promoters (HCP) bearing the H3K27 tri-methylation (H3K27me3) mark in brain.** | MEISSNER_BRAIN_HCP_WITH_H3K27ME3 | 2 | **4.6380e-9** | **1.5602e-5** | **7.8010e-6** | 3.4301 | 9.0376 | 31 | 0.0233 | 7.99% | 6 | **5.7355e-8** | **1.9294e-4** | **3.2157e-5** | 3.3023 | [illegible] |
| **Genes up-regulated in B-CLL (B-cell chronic lymphocytic leukemia) patients with mutated immunoglobulin variable heavy chain (VH) genes.** | FAELT_B_CLL_WITH_VH_REARRANGEMENTS_UP | 3 | **3.4497e-8** | **1.1605e-4** | **3.8683e-5** | 7.5167 | 1.7295 | 13 | 0.0045 | 3.35% | 14 | **1.8171e-6** | **6.1126e-3** | **4.3661e-4** | 6.6759 | [illegible] |
| **Genes up-regulated in acute myeloid leukemia (AML) with respect to cellular localization of NPM1 [GeneID=4869]: cytoplasmic vs. nucleolar.** | ALCALAY_AML_BY_NPM1_LOCALIZATION_UP | 4 | **1.6144e-7** | **5.4308e-4** | **1.3577e-4** | 3.5283 | 6.8021 | 24 | 0.0175 | 6.19% | 10 | **3.3546e-7** | **1.1285e-3** | **1.1285e-4** | 4.1496 | [illegible] |
| **Genes within amplicon 7p15 identified in a copy number alterations study of 191 breast tumor samples.** | NIKOLSKY_BREAST_CANCER_7P15_AMPLICON | 5 | **2.3136e-7** | **7.7830e-4** | **1.5566e-4** | 81.2873 | 0.0492 | 4 | 0.0001 | 1.03% | 49 | **2.5997e-4** | **8.7454e-1** | **1.7848e-2** | 11.6525 | [illegible] |
| **Top genes associated with unfavorable overall survival of mesothelioma patients after surgery.** | LOPEZ_MESOTHELIOMA_SURVIVAL_OVERALL_DN | 6 | **4.5845e-7** | **1.5422e-3** | **2.5704e-4** | 15.6514 | 0.4472 | 7 | 0.0012 | 1.80% | 79 | **9.7346e-4** | **1.0000** | **4.1452e-2** | 8.5452 | [illegible] |
| **The 'NPM1-mutated signature 2': genes down-regulated in pediatric AML (acute myeloid leukemia) samples with mutated NPM1 [GeneID=4869] compared to the AML cases with the intact gene and without recurring cytogenetic anomalities or M7 phenotype.** | MULLIGHAN_NPM1_MUTATED_SIGNATURE_2_DN | 7 | **3.1798e-6** | **1.0697e-2** | **1.5281e-3** | 5.4549 | 2.1999 | 12 | 0.0057 | 3.09% | 72 | **6.4316e-4** | **1.0000** | **3.0050e-2** | 3.7454 | [illegible] |
| **Genes down-regulated in normal hematopoietic progenitors by RUNX1-RUNX1T1 [GeneID=861;862] fusion.** | TONKS_TARGETS_OF_RUNX1_RUNX1T1_FUSION_HSC_DN | 8 | **3.7644e-6** | **1.2663e-2** | **1.5829e-3** | 3.2295 | 6.5026 | 21 | 0.0168 | 5.41% | 20 | **1.3088e-5** | **4.4029e-2** | **2.2014e-3** | 3.2223 | [illegible] |
| **The 'NPM1 signature 3': genes down-regulated in pediatric AML (acute myeloid leukemia) with mutated NPM1 [GeneID=4869] compared to the AML cases with intact NPM1 and MLL [GeneID=4297].** | MULLIGHAN_NPM1_SIGNATURE_3_DN | 9 | **6.1446e-6** | **2.0670e-2** | **2.2967e-3** | 3.2386 | 6.1755 | 20 | 0.0159 | 5.15% | 22 | **1.4035e-5** | **4.7212e-2** | **2.1460e-3** | 3.3421 | [illegible] |
| **Genes down-regulated in H358 cells (lung cancer) by inducible expression of CEBPA [GeneID=1050] off plasmid vector.** | HALMOS_CEBPA_TARGETS_DN | 10 | **7.8105e-6** | **2.6274e-2** | **2.6274e-3** | 4.9842 | 2.4076 | 12 | 0.0062 | 3.09% | 19 | **1.2445e-5** | **4.1867e-2** | **2.2035e-3** | 6.1362 | [illegible] |
| **Genes that best predicted acute myeloid leukemia (AML) with internal tandem duplications (IDT) in FLT3 [GeneID=2322].** | VALK_AML_WITH_FLT3_ITD | 12 | **1.0102e-5** | **3.3982e-2** | **2.8319e-3** | 5.9444 | 1.6823 | 10 | 0.0043 | 2.58% | 41 | **1.5107e-4** | **5.0820e-1** | **1.2395e-2** | 5.9029 | [illegible] |
| **The 'NPM1-mutated signature 1': genes down-regulated in pediatric AML (acute myeloid leukemia) samples with mutated NPM1 [GeneID=4869] compared to all AML cases with the intact gene.** | MULLIGHAN_NPM1_MUTATED_SIGNATURE_1_DN | 15 | **2.0587e-5** | **6.9254e-2** | **4.6169e-3** | 3.5028 | 4.5678 | 16 | 0.0118 | 4.12% | 43 | **1.7091e-4** | **5.7493e-1** | **1.3370e-2** | 3.2801 | [illegible] |
| **The 'NPM1-mutated signature 1': genes up-regulated in pediatric AML (acute myeloid leukemia) samples with mutated NPM1 [GeneID=4869] compared to all AML cases with the intact gene.** | MULLIGHAN_NPM1_MUTATED_SIGNATURE_1_UP | 16 | **2.4882e-5** | **8.3703e-2** | **5.2314e-3** | 2.4457 | 11.0399 | 27 | 0.0285 | 6.96% | 16 | **4.8181e-6** | **1.6208e-2** | **1.0130e-3** | 2.7624 | [illegible] |
| **Genes up-regulated during pubertal mammary gland development between week 4 and 5.** | MCBRYAN_PUBERTAL_BREAST_4_5WK_UP | 18 | **3.4718e-5** | **1.1679e-1** | **6.4883e-3** | 2.3099 | 12.5549 | 29 | 0.0324 | 7.47% | 9 | **2.4832e-7** | **8.3534e-4** | **9.2816e-5** | 3.1559 | [illegible] |
| **Genes up-regulated in normal bone marrow plasma cells (BMPC) compared to polyclonal plasmablasts (PPC) that also distinguished multiple myeloma (MM) samples by expression of levels of TACI (TNFRSF13B) [GeneID=23495].** | MOREAUX_B_LYMPHOCYTE_MATURATION_BY_TACI_UP | 19 | **4.5140e-5** | **1.5185e-1** | **7.9922e-3** | 4.5139 | 2.4369 | 11 | 0.0063 | 2.84% | 87 | **1.1114e-3** | **1.0000** | **4.2972e-2** | 3.4747 | [illegible] |
| **Genes down-regulated in mARMS (molecular ARMS) compared to the mERMS (molecular ERMS) class of rhabdomyosarcoma tumors.** | DAVICIONI_MOLECULAR_ARMS_VS_ERMS_DN | 21 | **4.9509e-5** | **1.6655e-1** | **7.9309e-3** | 2.5022 | 9.5914 | 24 | 0.0247 | 6.19% | 7 | **1.0992e-7** | **3.6978e-4** | **5.2826e-5** | 3.8898 | [illegible] |

| Name | ID | Rank | P-Value | P-Value | *Val* | Enrichment | Expected | Region Hits | Fraction | Set Coverage | Rank | P-Value | P-Value | *Val* | Enrichment | Ex |
|---|---|---|---|---|---|---|---|---|---|---|---|---|---|---|---|---|
| Genes up-regulated in SaOS-2 cells (osteosarcoma) upon knockdown of YY1 [GeneID=7528] by RNAi. | DE_YY1_TARGETS_UP | 22 | 6.0863e-5 | 2.0474e-1 | 9.3065e-3 | 9.1607 | 0.6550 | 6 | 0.0017 | 1.55% | 62 | 4.0794e-4 | 1.0000 | 2.2134e-2 | 6.0083 | ( |
| Genes down-regulated in HCT8/S11 cells (colon cancer) engineered to stably express NTN1 [GeneID=1630] off a plasmid vector. | RODRIGUES_NTN1_TARGETS_DN | 23 | 9.1648e-5 | 3.0830e-1 | 1.3405e-2 | 2.9553 | 5.7523 | 17 | 0.0148 | 4.38% | 63 | 4.1338e-4 | 1.0000 | 2.2073e-2 | 2.8575 | 4 |
| Genes with DNA sequences bound by RARA and RARG [GeneID=5914, 5916] in ES cells. | DELACROIX_RAR_BOUND_ES | 25 | 1.1679e-4 | 3.9287e-1 | 1.5715e-2 | 2.0883 | 14.8446 | 31 | 0.0383 | 7.99% | 50 | 2.6753e-4 | 8.9995e-1 | 1.7999e-2 | 2.0579 | 1 |
| Genes repressed in the time interval between two pulses of EGF [GeneID =1950] in 184A1 cells (mammary epithelium). | ZWANG_EGF_INTERVAL_DN | 26 | 1.2214e-4 | 4.1088e-1 | 1.5803e-2 | 2.5347 | 8.2849 | 21 | 0.0214 | 5.41% | 27 | 4.1549e-5 | 1.3977e-1 | 5.1767e-3 | 2.9579 | 6 |

The test set of 388 genomic regions picked 563 (3%) of all 18,041 genes.
*MSigDB Perturbation* has 3,364 terms covering 17,091 (95%) of all 18,041 genes, and 358,104 term - gene associations.
3,364 ontology terms (100%) were tested using an annotation count range of [1, Inf].

### MSigDB Predicted Promoter Motifs (20+ terms)

Global controls

*Table controls:* Export ▾ Shown top rows in this table: 20 Set Term annotation count: Min: 1 Max: Inf Set Visualize this table: NEW [select one] ▾

| Term Name | Term ID | Binom Rank | Binom Raw P-Value | Binom Bonferroni P-Value | *Binom FDR Q-Val* | Binom Fold Enrichment | Binom Expected | Binom Observed Region Hits | Binom Genome Fraction | Binom Region Set Coverage | Hyper Rank | Hyper Raw P-Value | Hyper Bonferroni P-Value | *Hyper FDR Q-Val* | Hyper Fold Enrichment | Hyper Expected | Hyper Observed Gene Hits | Hyper Total Genes | G C |
|---|---|---|---|---|---|---|---|---|---|---|---|---|---|---|---|---|---|---|---|
| Motif CANCCNNWGGGTGDGG (no known TF) | V$CACCCBINDINGFACTOR_Q6 | 2 | 1.1429e-7 | 7.0285e-5 | 3.5143e-5 | 2.7923 | 12.1763 | 34 | 0.0314 | 8.76% | 9 | 1.1461e-8 | 7.0488e-6 | 7.8320e-7 | 3.4643 | 8.0825 | 28 | 259 | |
| Motif GGCGSG matches E2F<br> TFDP1: transcription factor Dp-1 | V$E2F_Q2 | 3 | 6.5969e-7 | 4.0571e-4 | 1.3524e-4 | 3.4821 | 6.3180 | 22 | 0.0163 | 5.67% | 10 | 5.9458e-8 | 3.6567e-5 | 3.6567e-6 | 4.0295 | 5.2115 | 21 | 167 | |
| Motif GRGGSTGGG (no known TF) | V$CACBINDINGPROTEIN_Q6 | 4 | 2.5596e-6 | 1.5741e-3 | 3.9354e-4 | 3.0083 | 7.9779 | 24 | 0.0206 | 6.19% | 73 | 1.2927e-4 | 7.9502e-2 | 1.0891e-3 | 2.6243 | 7.2400 | 19 | 232 | |
| Motif NGNGGGGA (no known TF) | V$MZF1_01 | 5 | 3.4831e-6 | 2.1421e-3 | 4.2842e-4 | 2.7356 | 9.8700 | 27 | 0.0254 | 6.96% | 40 | 1.2302e-5 | 7.5659e-3 | 1.8915e-4 | 2.9131 | 7.2087 | 21 | 231 | |
| Motif NNGKNTGTGGTTWNC matches RUNX1: runt-related transcription factor 1 (acute myeloid leukemia 1; aml1 oncogene) | V$AML_Q6 | 6 | 5.7329e-6 | 3.5258e-3 | 5.8763e-4 | 2.5503 | 11.3711 | 29 | 0.0293 | 7.47% | 33 | 6.6255e-6 | 4.0747e-3 | 1.2347e-4 | 2.8567 | 8.0513 | 23 | 258 | |
| Motif GCHCDAMCCAG matches TFCP2: transcription factor CP2 | V$CP2_01 | 7 | 5.8144e-6 | 3.5758e-3 | 5.1083e-4 | 2.6013 | 10.7639 | 28 | 0.0277 | 7.22% | 31 | 5.8231e-6 | 3.5812e-3 | 1.1552e-4 | 2.8790 | 7.9889 | 23 | 256 | |
| Motif NNGTTGTTTACNTN matches MLLT7: myeloid/lymphoid or mixed-lineage leukemia (trithorax homolog, Drosophila); translocated to, 7 | V$FOXO4_02 | 8 | 7.8915e-6 | 4.8533e-3 | 6.0666e-4 | 2.2352 | 16.1061 | 36 | 0.0415 | 9.28% | 5 | 1.4785e-9 | 9.0927e-7 | 1.8185e-7 | 3.6877 | 7.8641 | 29 | 252 | |
| Motif NNTTTCCN matches STAT1: signal transducer and activator of transcription 1, 91kDa | V$STAT1_03 | 9 | 1.0086e-5 | 6.2030e-3 | 6.8922e-4 | 2.6986 | 9.2640 | 25 | 0.0239 | 6.44% | 103 | 5.8812e-4 | 3.6169e-1 | 3.5116e-3 | 2.3934 | 7.5208 | 18 | 241 | |
| Motif GCGSCMNTTT (no known TF) | GCGSCMNTTT_UNKNOWN | 10 | 1.8928e-5 | 1.1641e-2 | 1.1641e-3 | 4.5524 | 2.6360 | 12 | 0.0068 | 3.09% | 179 | 4.7865e-3 | 1.0000 | 1.6445e-2 | 3.3479 | 2.0908 | 7 | 67 | |
| Motif CNGNRNCAGGTGNNGNAN matches MYOD1: myogenic differentiation 1 | V$MYOD_Q6_01 | 11 | 2.0476e-5 | 1.2592e-2 | 1.1448e-3 | 2.5269 | 10.2891 | 26 | 0.0265 | 6.70% | 61 | 7.2169e-5 | 4.4384e-2 | 7.2761e-4 | 2.6593 | 7.5208 | 20 | 241 | |
| Motif ATGCCCATATATGGWNNT matches SRF: serum response factor (c-fos serum response element-binding transcription factor) | V$SRF_01 | 14 | 9.1845e-5 | 5.6484e-2 | 4.0346e-3 | 5.7981 | 1.3798 | 8 | 0.0036 | 2.06% | 165 | 4.0255e-3 | 1.0000 | 1.5004e-2 | 3.9238 | 1.5291 | 6 | 49 | |
| Motif CACSCCA matches SREBF1: sterol regulatory element binding transcription factor 1 | V$SREBP1_Q6 | 15 | 1.0584e-4 | 6.5093e-2 | 4.3396e-3 | 2.4345 | 9.4477 | 23 | 0.0243 | 5.93% | 79 | 1.6155e-4 | 9.9352e-2 | 1.2576e-3 | 2.5798 | 7.3648 | 19 | 236 | |

| Name | ID | Rank | P-Value | Bonferroni P-Value | FDR Q-Val | Fold Enrichment | Expected | Region Hits | Genome Fraction | Set Coverage | Rank | Raw P-Value | Bonferroni P-Value | FDR Q-Val | Fold Enrichment | Expected | Gene Hits | Total Genes | C |
|---|---|---|---|---|---|---|---|---|---|---|---|---|---|---|---|---|---|---|---|
| **Motif TATAAATW matches TBP: TATA box binding protein** | V$TBP_01 | 16 | **1.1421e-4** | **7.0239e-2** | **4.3899e-3** | 2.2304 | 12.1054 | 27 | 0.0312 | 6.96% | 70 | **1.2215e-4** | **7.5123e-2** | **1.0732e-3** | 2.6357 | 7.2087 | 19 | 231 | |
| **Motif TGTGGTTW matches CBFA2T2: core-binding factor, runt domain, alpha subunit 2; translocated to, 2<br> CBFA2T3: core-binding factor, runt domain, alpha subunit 2; translocated to, 3** | V$COREBINDINGFACTOR_Q6 | 17 | **1.2123e-4** | **7.4556e-2** | **4.3857e-3** | 2.2221 | 12.1505 | 27 | 0.0313 | 6.96% | 107 | **7.5573e-4** | **4.6477e-1** | **4.3437e-3** | 2.2803 | 8.3322 | 19 | 267 | |
| **Motif TGACCTTTGNCCY matches PPARA: peroxisome proliferative activated receptor, alpha** | V$PPAR_DR1_Q2 | 18 | **1.6267e-4** | **1.0004e-1** | **5.5580e-3** | 2.4813 | 8.4632 | 21 | 0.0218 | 5.41% | 239 | **1.1379e-2** | **1.0000** | **2.9281e-2** | 1.9382 | 7.7393 | 15 | 248 | |
| **Motif NNNTTCCN matches STAT3: signal transducer and activator of transcription 3 (acute-phase response factor)** | V$STAT3_02 | 19 | **1.7277e-4** | **1.0625e-1** | **5.5922e-3** | 2.6153 | 7.2649 | 19 | 0.0187 | 4.90% | 83 | **1.9500e-4** | **1.1993e-1** | **1.4449e-3** | 3.0728 | 4.5562 | 14 | 146 | |
| **Motif NNNGGNCNCAGCTGCGNCCCNN matches NHLH1: nescient helix loop helix 1** | V$HEN1_01 | 20 | **1.8748e-4** | **1.1530e-1** | **5.7649e-3** | 2.3942 | 9.1888 | 22 | 0.0237 | 5.67% | 42 | **1.3799e-5** | **8.4861e-3** | **2.0205e-4** | 3.0906 | 6.1477 | 19 | 197 | |
| **Motif NNNRRCCAATSRGNNN (no known TF)** | V$NFY_01 | 22 | **1.8885e-4** | **1.1614e-1** | **5.2792e-3** | 2.2877 | 10.4907 | 24 | 0.0270 | 6.19% | 138 | **1.9214e-3** | **1.0000** | **8.5628e-3** | 2.2145 | 7.6768 | 17 | 246 | |
| **Motif CCCCAAACMMCCCC matches RREB1: ras responsive element binding protein 1** | V$RREB1_01 | 23 | **2.3667e-4** | **1.4555e-1** | **6.3285e-3** | 2.3537 | 9.3471 | 22 | 0.0241 | 5.67% | 60 | **5.7951e-5** | **3.5640e-2** | **5.9400e-4** | 2.8840 | 6.2413 | 18 | 200 | |
| **Motif NNNTGGGAWNNC (no known TF)** | V$IK2_01 | 24 | **2.3709e-4** | **1.4581e-1** | **6.0755e-3** | 2.0326 | 14.7596 | 30 | 0.0380 | 7.73% | 24 | **2.2839e-6** | **1.4046e-3** | **5.8524e-5** | 2.9579 | 8.1137 | 24 | 260 | |

The test set of 388 genomic regions picked 563 (3%) of all 18,041 genes.
*MSigDB Predicted Promoter Motifs* has 615 terms covering 11,991 (66%) of all 18,041 genes, and 158,271 term - gene associations.
615 ontology terms (100%) were tested using an annotation count range of [1, Inf].

### MSigDB miRNA Motifs (no terms)

Global controls

*Table controls:* Export ▾ Shown top rows in this table: 20 Set Term annotation count: Min: 1 Max: Inf Set Visualize this table: [select one] ▾

| Term Name | Term ID | Binom Rank | Binom Raw P-Value | Binom Bonferroni P-Value | *Binom FDR Q-Val* | Binom Fold Enrichment | Binom Expected | Binom Observed Region Hits | Binom Genome Fraction | Binom Region Set Coverage | Hyper Rank | Hyper Raw P-Value | Hyper Bonferroni P-Value | *Hyper FDR Q-Val* | Hyper Fold Enrichment | Hyper Expected | Hyper Observed Gene Hits | Hyper Total Genes | Hyper Gene Set Coverage | Hyper Term Gene Coverage |
|---|---|---|---|---|---|---|---|---|---|---|---|---|---|---|---|---|---|---|---|---|

No results meet your chosen criteria.

The test set of 388 genomic regions picked 563 (3%) of all 18,041 genes.
*MSigDB miRNA Motifs* has 221 terms covering 7,064 (39%) of all 18,041 genes, and 32,882 term - gene associations.
221 ontology terms (100%) were tested using an annotation count range of [1, Inf].

### InterPro (6 terms)

Global controls

*Table controls:* Export ▾ Shown top rows in this table: 20 Set Term annotation count: Min: 1 Max: Inf Set Visualize this table: [select one] ▾

| Term Name | Term ID | Binom Rank | Binom Raw P-Value | Binom Bonferroni P-Value | *Binom FDR Q-Val* | Binom Fold Enrichment | Binom Expected | Binom Observed Region Hits | Binom Genome Fraction | Binom Region Set Coverage | Hyper Rank | Hyper Raw P-Value | Hyper Bonferroni P-Value | *Hyper FDR Q-Val* | Hyper Fold Enrichment | Hyper Expected | Hyper Observed Gene Hits | Hyper Total Genes | Hyper Gene Set Coverage | Hyper Term Gene Coverage |
|---|---|---|---|---|---|---|---|---|---|---|---|---|---|---|---|---|---|---|---|---|
| **Homeobox protein, antennapedia type, conserved site** | IPR001827 | 1 | **4.4089e-27** | **4.1541e-23** | **4.1541e-23** | 239.5964 | 0.0543 | 13 | 0.0001 | 3.35% | 5 | **1.1286e-10** | **1.0634e-6** | **2.1268e-7** | 16.0222 | 0.6241 | 10 | 20 | 1.78% | 50.00% |
| **Homeobox protein, antennapedia type** | IPR017995 | 2 | **1.3625e-19** | **1.2838e-15** | **6.4189e-16** | 167.1899 | 0.0598 | 10 | 0.0002 | 2.58% | 6 | **1.9197e-5** | **1.8088e-1** | **3.0146e-2** | 13.3518 | 0.3745 | 5 | 12 | 0.89% | 41.67% |

| Name | ID | Rank | P-Value | Bonferroni P-Value | FDR Q Val | Fold Enrichment | Expected | Region Hits | Genome Fraction | Set Coverage | Rank | Raw P-Value | Bonferroni P-Value | FDR Q Val | Fold Enrichment | Expected | Gene Hits | Total Genes | Gene Set Coverage | Gene Coverage |
|---|---|---|---|---|---|---|---|---|---|---|---|---|---|---|---|---|---|---|---|---|
| **Homeodomain, metazoa** | IPR020479 | 3 | **3.5144e-14** | **3.3112e-10** | **1.1037e-10** | 8.1590 | 2.8190 | 23 | 0.0073 | 5.93% | 4 | **5.6631e-12** | **5.3358e-8** | **1.3339e-8** | 6.9662 | 2.8710 | 20 | 92 | 3.55% | 21.74% |
| **Homeobox, conserved site** | IPR017970 | 4 | **3.7423e-11** | **3.5260e-7** | **8.8149e-8** | 3.8671 | 8.7921 | 34 | 0.0227 | 8.76% | 1 | **2.6966e-14** | **2.5408e-10** | **2.5408e-10** | 5.2839 | 5.8669 | 31 | 188 | 5.51% | 16.49% |
| **Homeobox domain** | IPR001356 | 5 | **2.2310e-9** | **2.1020e-5** | **4.2041e-6** | 3.0338 | 12.5253 | 38 | 0.0323 | 9.79% | 2 | **3.8739e-14** | **3.6500e-10** | **1.8250e-10** | 4.6154 | 7.5832 | 35 | 243 | 6.22% | 14.40% |
| **Homeodomain-like** | IPR009057 | 6 | **2.4742e-8** | **2.3312e-4** | **3.8853e-5** | 2.6000 | 16.1537 | 42 | 0.0416 | 10.82% | 3 | **6.6471e-13** | **6.2629e-9** | **2.0876e-9** | 3.8218 | 10.2046 | 39 | 327 | 6.93% | 11.93% |

The test set of 388 genomic regions picked 563 (3%) of all 18,041 genes.
*InterPro* has 9,422 terms covering 17,343 (96%) of all 18,041 genes, and 60,477 term - gene associations.
9,422 ontology terms (100%) were tested using an annotation count range of [1, Inf].

### TreeFam (no terms)

Global controls

*Table controls:* Export ▾ Shown top rows in this table: 20 Set Term annotation count: Min: 1 Max: Inf Set Visualize this table: NEW [select one] ▾

| Term Name | Term ID | Binom Rank | Binom Raw P-Value | Binom Bonferroni P-Value | *Binom FDR Q-Val* | Binom Fold Enrichment | Binom Expected | Binom Observed Region Hits | Binom Genome Fraction | Binom Region Set Coverage | Hyper Rank | Hyper Raw P-Value | Hyper Bonferroni P-Value | *Hyper FDR Q-Val* | Hyper Fold Enrichment | Hyper Expected | Hyper Observed Gene Hits | Hyper Total Genes | Hyper Gene Set Coverage | Hyper Term Gene Coverage |
|---|---|---|---|---|---|---|---|---|---|---|---|---|---|---|---|---|---|---|---|---|

No results meet your chosen criteria.

The test set of 388 genomic regions picked 563 (3%) of all 18,041 genes.
*TreeFam* has 8,126 terms covering 13,550 (75%) of all 18,041 genes, and 13,551 term - gene associations.
8,126 ontology terms (100%) were tested using an annotation count range of [1, Inf].

### HGNC Gene Families (1 term)

Global controls

*Table controls:* Export ▾ Shown top rows in this table: 20 Set Term annotation count: Min: 1 Max: Inf Set Visualize this table: NEW [select one] ▾

| Term Name | Term ID | Binom Rank | Binom Raw P-Value | Binom Bonferroni P-Value | *Binom FDR Q-Val* | Binom Fold Enrichment | Binom Expected | Binom Observed Region Hits | Binom Genome Fraction | Binom Region Set Coverage | Hyper Rank | Hyper Raw P-Value | Hyper Bonferroni P-Value | *Hyper FDR Q-Val* | Hyper Fold Enrichment | Hyper Expected | Hyper Observed Gene Hits | Hyper Total Genes | Hyper Gene Set Coverage | Hyper Term Gene Coverage |
|---|---|---|---|---|---|---|---|---|---|---|---|---|---|---|---|---|---|---|---|---|
| **HOXL** | HOXL | 1 | **1.5139e-18** | **7.2366e-16** | **7.2366e-16** | 25.0014 | 0.6800 | 17 | 0.0018 | 4.38% | 1 | **3.3259e-11** | **1.5898e-8** | **1.5898e-8** | 9.2436 | 1.6227 | 15 | 52 | 2.66% | 28.85% |

The test set of 388 genomic regions picked 563 (3%) of all 18,041 genes.
*HGNC Gene Families* has 478 terms covering 7,310 (41%) of all 18,041 genes, and 7,310 term - gene associations.
478 ontology terms (100%) were tested using an annotation count range of [1, Inf].

### Ensembl Genes (no terms)

Global controls

*Table controls:* Export ▾ Shown top rows in this table: 20 Set Term annotation count: Min: 1 Max: Inf Set Visualize this table: NEW [select one] ▾

| Term Name | Term ID | Binom Rank | Binom Raw P-Value | Binom Bonferroni P-Value | *Binom FDR Q-Val* | Binom Fold Enrichment | Binom Expected | Binom Observed Region Hits | Binom Genome Fraction | Binom Region Set Coverage | Hyper Rank | Hyper Raw P-Value | Hyper Bonferroni P-Value | *Hyper FDR Q-Val* | Hyper Fold Enrichment | Hyper Expected | Hyper Observed Gene Hits | Hyper Total Genes | Hyper Gene Set Coverage | Hyper Term Gene Coverage |
|---|---|---|---|---|---|---|---|---|---|---|---|---|---|---|---|---|---|---|---|---|

No results meet your chosen criteria.

18,041 ontology terms (100%) were tested using an annotation count range of [1, Inf].

### MSigDB Oncogenic Signatures (2 terms)

Global controls

*Table controls:* Export ▾ Shown top rows in this table: 20 Set Term annotation count: Min: 1 Max: Inf Set Visualize this table: [select one] ▾

| Term Name | Term ID | Binom Rank | Binom Raw P-Value | Binom Bonferroni P-Value | *Binom FDR Q-Val* | Binom Fold Enrichment | Binom Expected | Binom Observed Region Hits | Binom Genome Fraction | Binom Region Set Coverage | Hyper Rank | Hyper Raw P-Value | Hyper Bonferroni P-Value | *Hyper FDR Q-Val* | Hyper Fold Enrichment | Hyper Expected | Hyper Observed Gene Hits | Hyper Total Genes | Hyper Gene Set Coverage | Hy Te Ge Cove |
|---|---|---|---|---|---|---|---|---|---|---|---|---|---|---|---|---|---|---|---|---|
| **Genes up-regulated in small intenstine in PRKCA [Gene ID=5578] knockout mice.** | PKCA_DN.V1_UP | 1 | **2.7178e-6** | **5.0823e-4** | **5.0823e-4** | 3.9160 | 4.3412 | 17 | 0.0112 | 4.38% | 6 | **1.8345e-3** | **3.4305e-1** | **5.7175e-2** | 2.5557 | 5.0867 | 13 | 163 | 2.31% | 7.9 |
| **Genes up-regulated in Sez-4 cells (T lymphocyte) that were first starved of IL2 [Gene ID=3558] and then stimulated with IL2 [Gene ID=3558].** | IL2_UP.V1_UP | 2 | **1.9864e-4** | **3.7145e-2** | **1.8573e-2** | 2.5861 | 7.3469 | 19 | 0.0189 | 4.90% | 3 | **5.0627e-4** | **9.4672e-2** | **3.1557e-2** | 2.6853 | 5.5860 | 15 | 179 | 2.66% | 8.3 |

The test set of 388 genomic regions picked 563 (3%) of all 18,041 genes.
*MSigDB Oncogenic Signatures* has 187 terms covering 10,315 (57%) of all 18,041 genes, and 30,004 term - gene associations.
187 ontology terms (100%) were tested using an annotation count range of [1, Inf].

### MSigDB Immunologic Signatures (no terms)

Global controls

*Table controls:* Export ▾ Shown top rows in this table: 20 Set Term annotation count: Min: 1 Max: Inf Set Visualize this table: [select one] ▾

| Term Name | Term ID | Binom Rank | Binom Raw P-Value | Binom Bonferroni P-Value | *Binom FDR Q-Val* | Binom Fold Enrichment | Binom Expected | Binom Observed Region Hits | Binom Genome Fraction | Binom Region Set Coverage | Hyper Rank | Hyper Raw P-Value | Hyper Bonferroni P-Value | *Hyper FDR Q-Val* | Hyper Fold Enrichment | Hyper Expected | Hyper Observed Gene Hits | Hyper Total Genes | Hyper Gene Set Coverage | Hyper Term Gene Coverage |
|---|---|---|---|---|---|---|---|---|---|---|---|---|---|---|---|---|---|---|---|---|

No results meet your chosen criteria.

The test set of 388 genomic regions picked 563 (3%) of all 18,041 genes.
*MSigDB Immunologic Signatures* has 1,910 terms covering 16,609 (92%) of all 18,041 genes, and 363,333 term - gene associations.
1,910 ontology terms (100%) were tested using an annotation count range of [1, Inf].

STANFORD SCHOOL OF MEDICINE Bejerano Lab STANFORD ENGINEERING

Copyright © 2010-2017. The Board of Trustees of Leland Stanford Junior University. All rights reserved.
